## Supplementary information for "3D Variability Analysis Reveals a Hidden Conformational Change Controlling Ammonia Transport in Human Asparagine Synthetase"

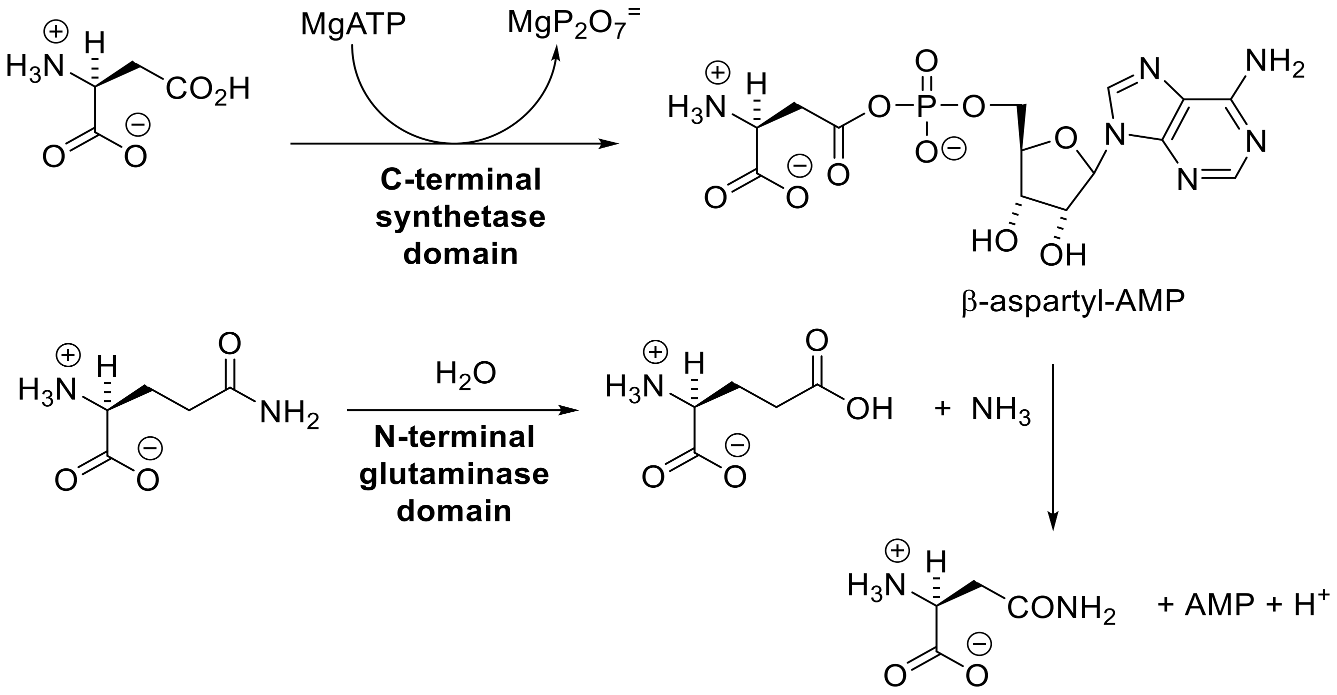

**Figure S1**. **Reactions catalyzed by the active sites in the N- and C-terminal domains of asparagine synthetase.** Ammonia released in the glutaminase site is channeled to the synthetase site through an intramolecular tunnel linking the two active sites.

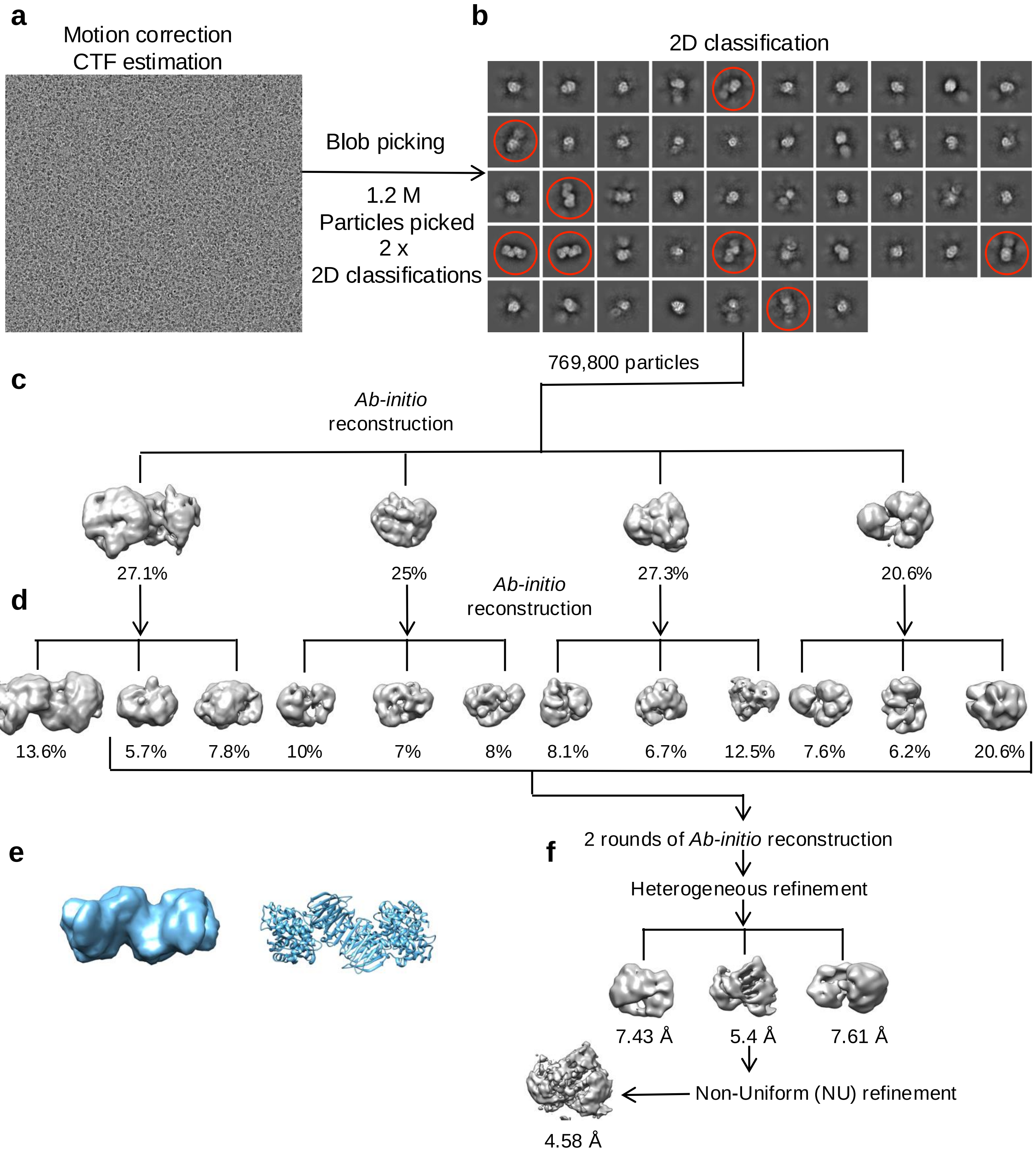

**Figure S2**. **Unbiased initial image analysis to assess the mode of ASNS dimerization.**

**a** Representative motion corrected micrograph.

**b** 2D class average: 2D projections resembling to the head-to-head dimer are indicated (red circle).

**c** The first round of Ab-initio reconstruction, yielding 4 different reconstructions.

**d** 2$^{nd}$ round of Ab-initio reconstruction, yielding a total of 12 different reconstructions.

**e** Stimulated EM density at 12 Å (on the left) derived from the crystal structure (PDB: 6GQ3) on the right.

**f** Heterogeneous refinement resulting in 3 reconstructions followed by NU refinement.

a

b

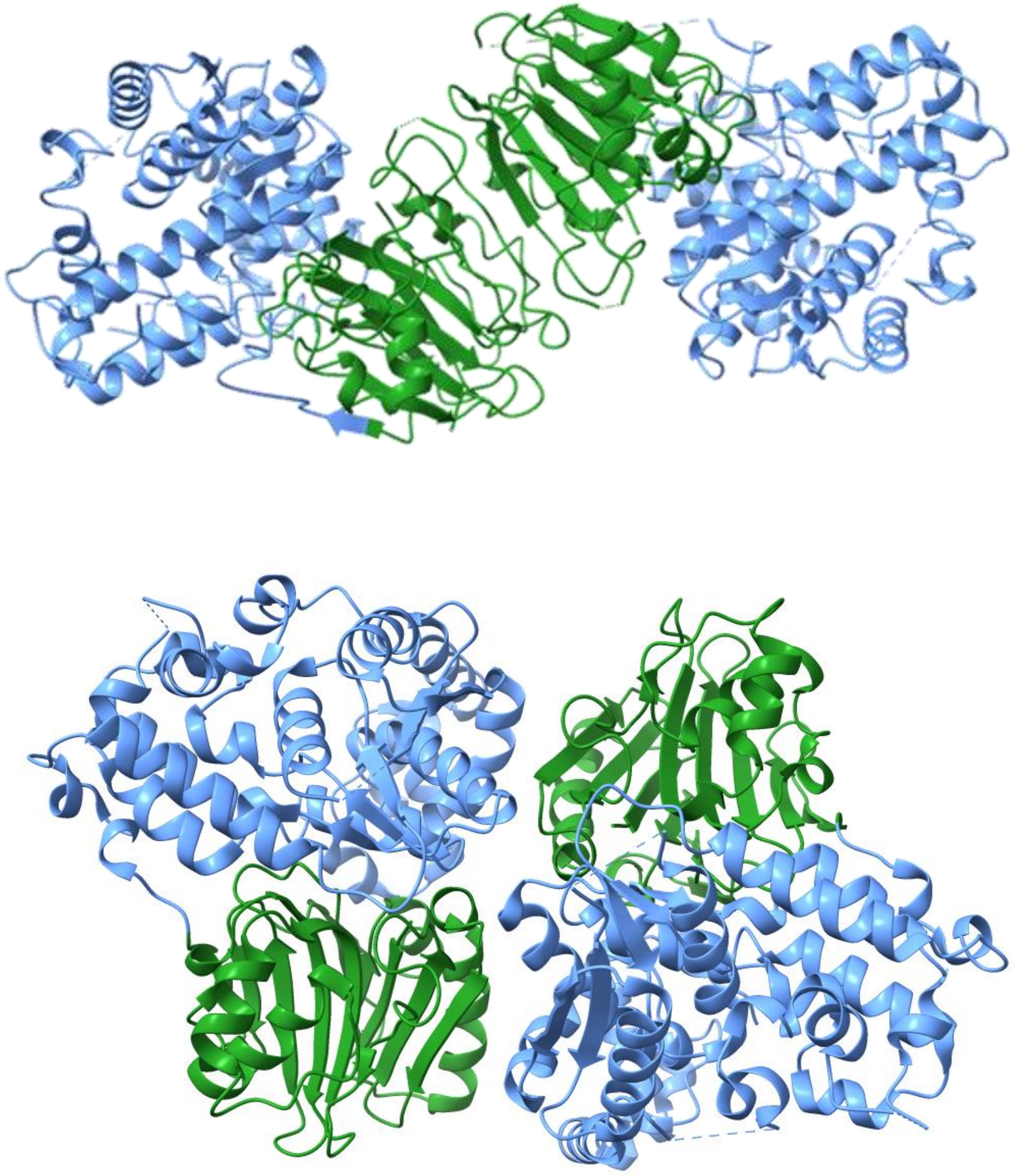

**Figure S3**. **Cartoon representations of the human and bacterial ASNS dimer. a** Ribbon diagram of the "head-to-head" dimer formed by human ASNS. **b.** Ribbon diagram of the "head-to-tail" dimer observed in the X-ray crystal structure of *Escherichia coli* AS-B. The N- and C-terminal domains of both structures are colored green and blue, respectively.

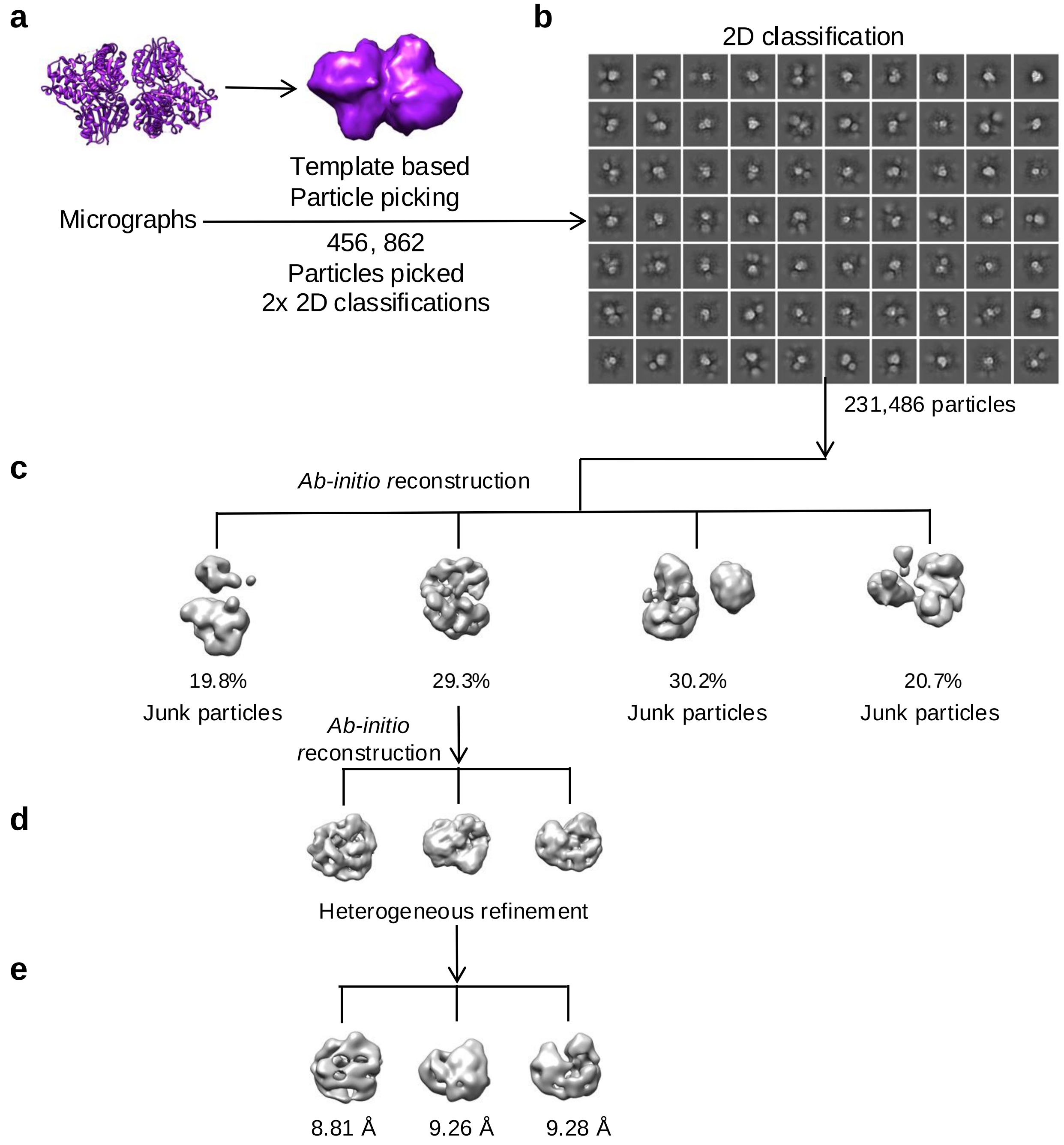

**Figure S4**. **Image analysis using AS-B as a template to assess the mode of ASNS dimerization.**
**a** Stimulated EM density at 12 Å (on the right) derived from the crystal structure (PDB: 1CT9) on the left.
**b** Selected 2D class averages of particles picked by templated-based approach.
**c** Ab-initio reconstruction, yielding 4 different reconstructions, 3 of which were considered to be junk.
**d** 2nd round of Ab-initio reconstruction, yielding a total of 3 reconstructions.
**e** Heterogeneous refinement resulting in a total of 3 3D reconstructions.

**Table S1. Cryo-EM data collection, refinement and validation statistics**

| **Data collection** | | |
|---|---|---|
| | apo-ASNS | ASNS(R142I) |
| Magnification | ×150,000 | X150,000 |
| Voltage (kV) | 200 | 200 |
| Electron exposure (e/Å$^2$) | 40 | 40 |
| Defocus range (μm) | -0.8 to -2.0 | -0.8 to -2.0 |
| Pixel size (Å) | 0.93 | 0.93 |
| **Data processing** | | |
| | **EMD-40764, PDB 8SUE** | **EMD-44253, PDB 9B6C** |
| Symmetry imposed | C2 | C2 |
| Initial particle images (no.) | 2,464,389 | 3,123,932 |
| Final particle images (no.) | 47,929 | 145,711 |
| Map resolution (Å) | 3.53 | 3.35 |
| FSC threshold | 0.143 | 0.143 |
| Map resolution range | 2.8-4.0 | 2.5-3.5 |
| Map sharpening B factor (Å$^2$) | 119.8 | 118.5 |
| **Refinement** | | |
| Model resolution (Å$^2$) | 2.3 | 2.2 |
| Model composition | | |
| Non-hydrogen atoms | 8049 | 8124 |
| Protein residues | 1019 | 1031 |
| Ligand (Mg) | 0 | 0 |
| B-factors (min/max/mean) | | |
| Protein | 36.67/74.22/51.65 | 10.29/103.04/36.04 |
| Ligand | --- | --- |
| r.m.s.d. deviations | | |
| Bond length (Å) | 0.003 | 0.004 |
| Bond angles (°) | 0.638 | 0.571 |
| Validation | | |
| MolProbility score | 2.05 | 1.91 |
| Clashscore | 12.53 | 7.78 |
| Rotamer outliers (%) | 0.36 | 1.65 |
| CaBLAM outliers (%) | 3.5 | 2.83 |
| Ramachandran plot (%) | | |
| Outliers | 0 | 0 |
| Allowed | 6.83 | 4.66 |
| Favored | 93.17 | 95.34 |

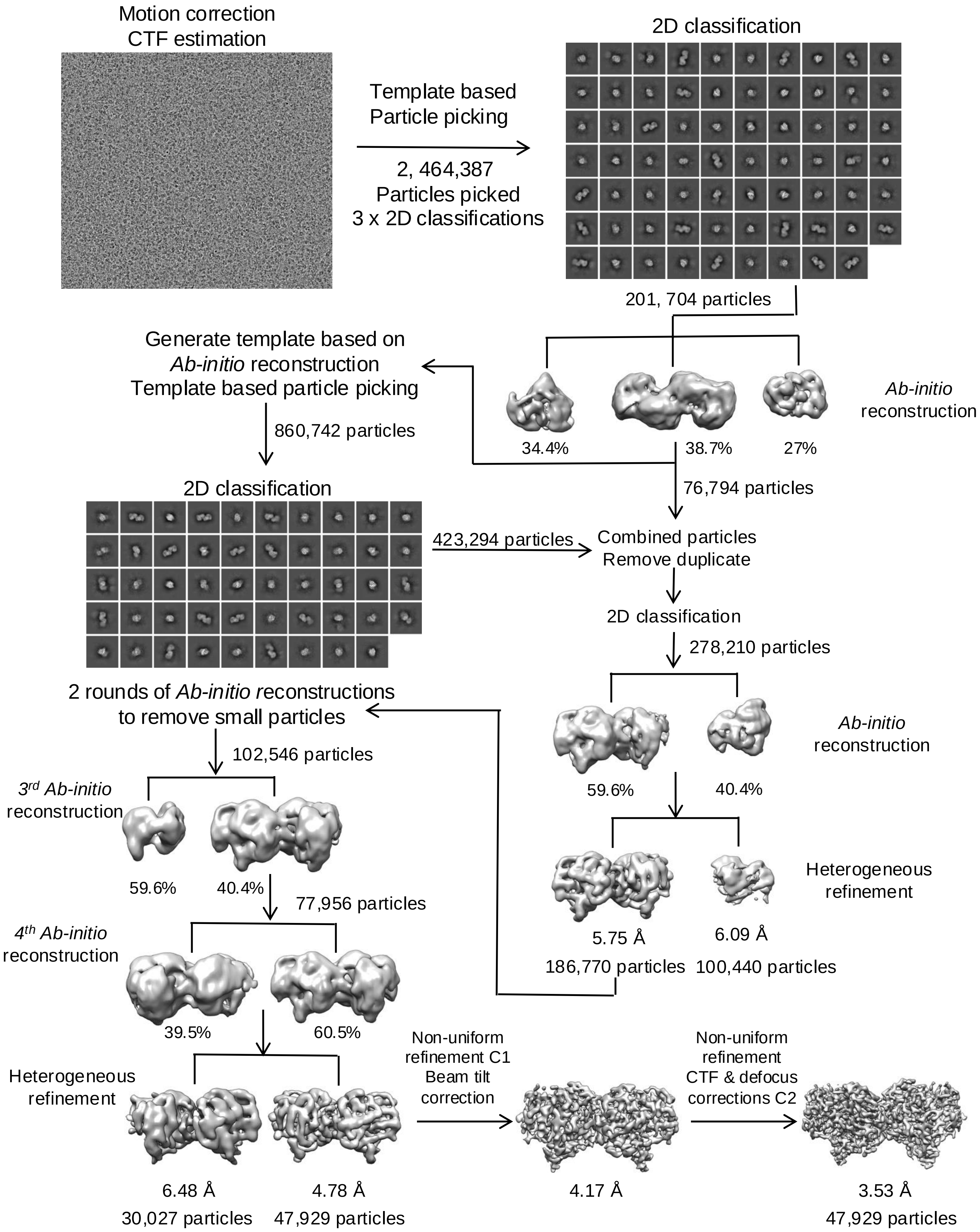

**Figure S5. Summary of the cryo-EM data processing workflow using cryoSPARC v3.2.2.**

**a Gold standard FSC curve for WT human ASNS**

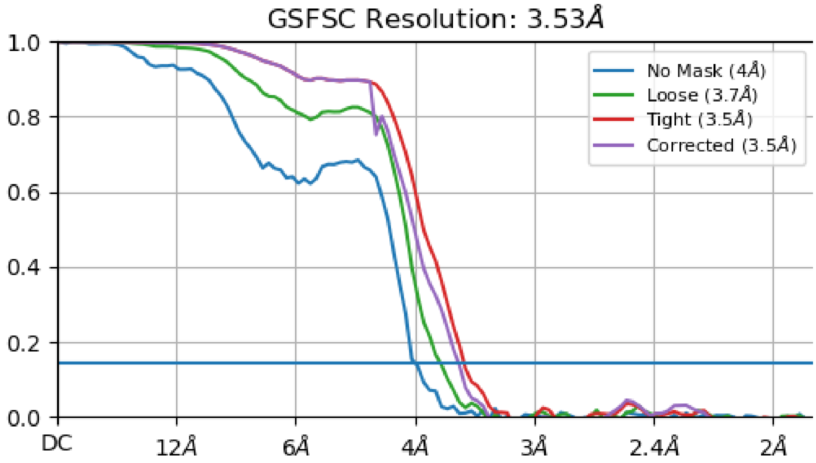

**b Viewing distribution plots**

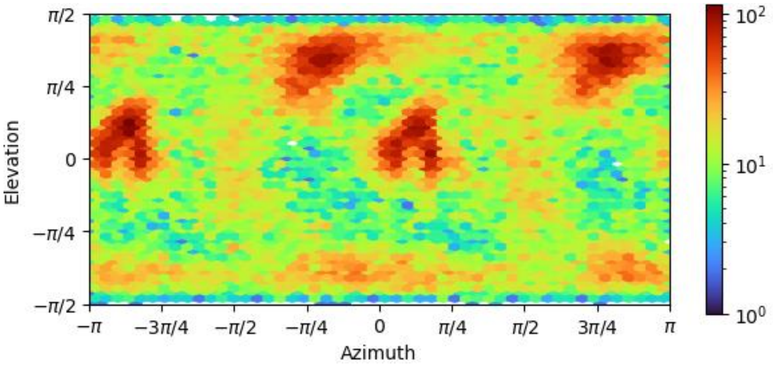

**c 3D FSC**

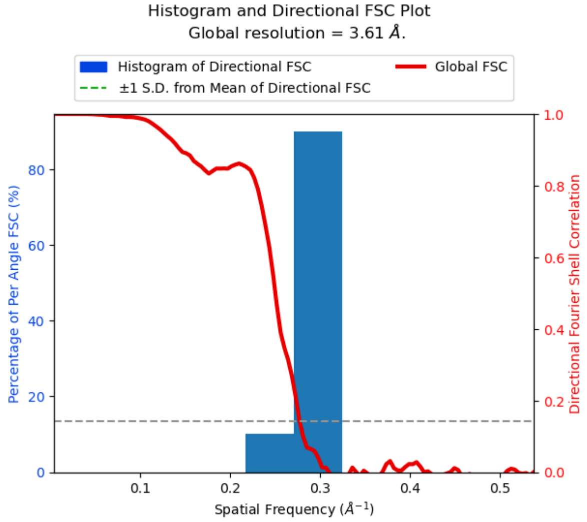

**d Local resolution**

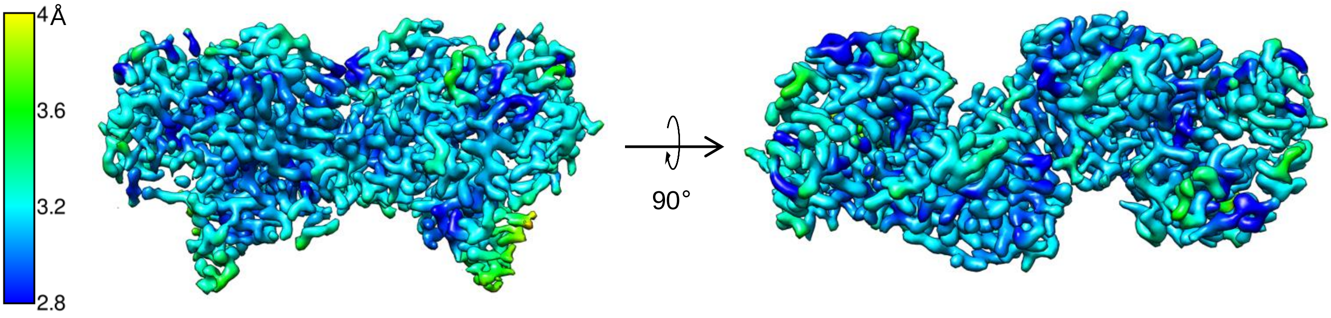

**Figure S6. Assessment of the quality of cryo-EM map.**
**a** Gold standard FSC curve for the map of hASNS generated in cryoSPARC v3.2.
**b** Viewing distribution plots for the map generated in cryoSPARC v3.2.
**c** 3D FSC: histogram and directional FSC plot for map global resolution are shown.
**d** Local resolution of the map (color scale shown on the left).

### a Representative densities and corresponding models for WT human ASNS

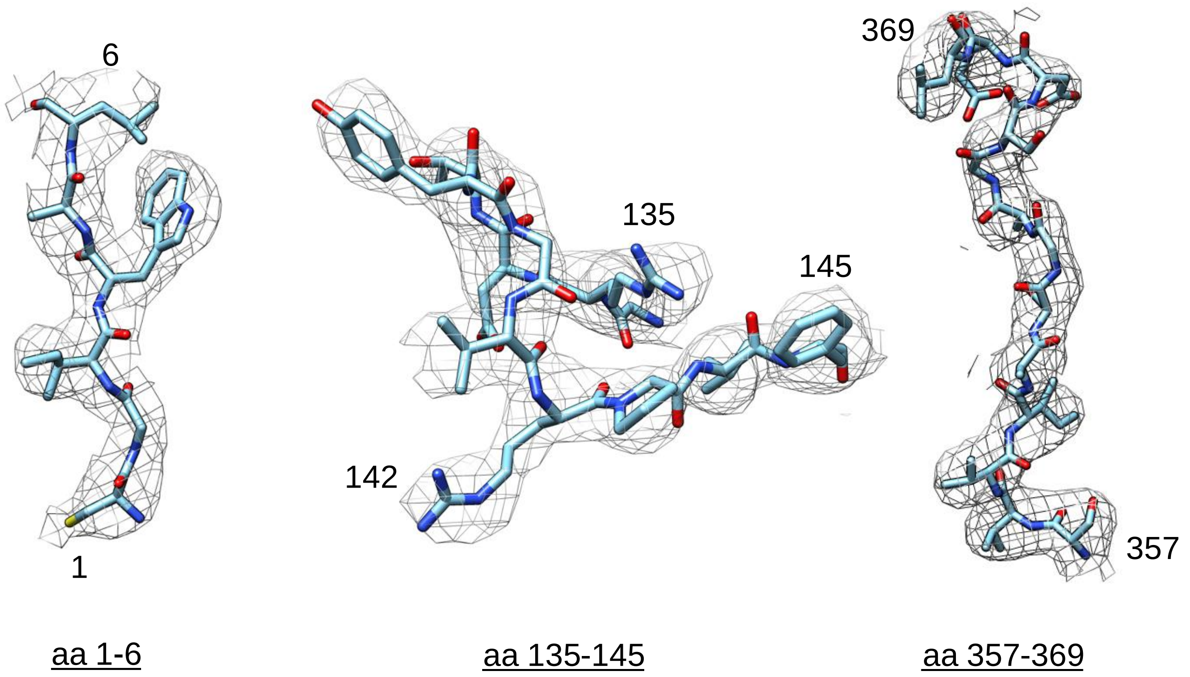

### b Map-model FSC curve

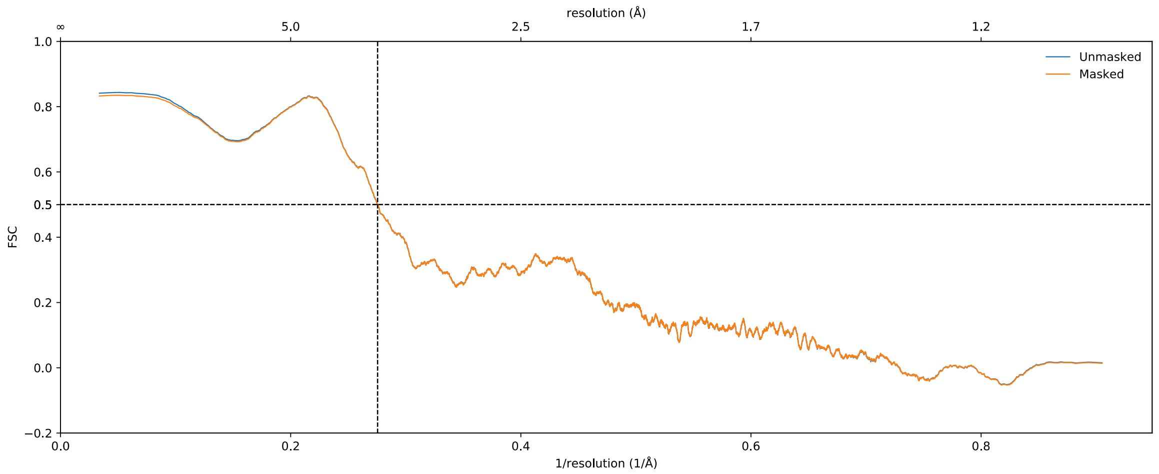

**Figure S7. Validating the cryo-EM structure: map and its model.**

**a** Representative density of the map for residues in the N-terminal catalytic region (aa 1-6), the ammonia tunnel region (aa 132-145), and the C-terminal active site (aa 357-369)

**b** Model-to-map FSC curves generated in Phenix.

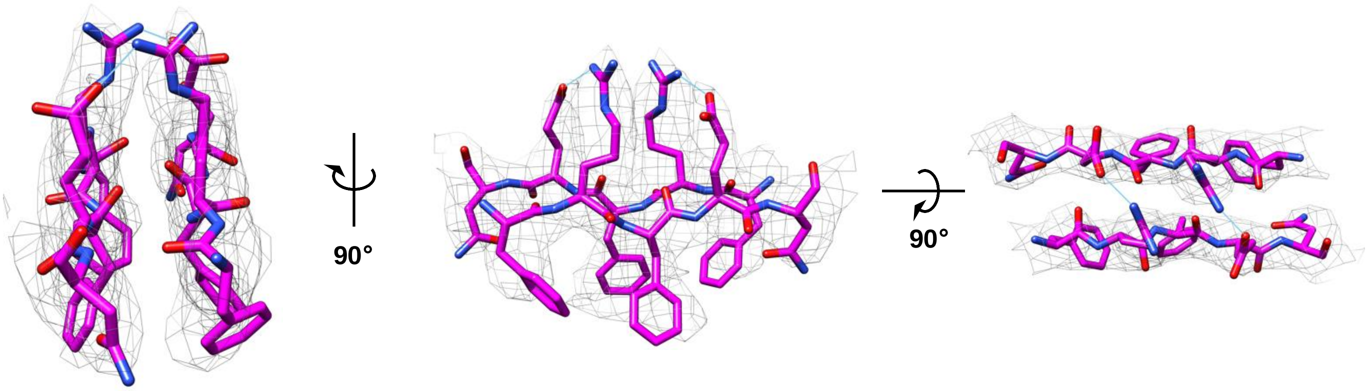

**Figure S8. Dimerization region with the map and model.** Density from the EM map is shown for residues located in the dimerization region (aa 30-35) is displayed with the model. Salt bridges in between residues Arg-32 or Glu-34 (monomer A) and Glu-34 or Arg-32 (monomer B) are indicated by lines colored in cyan. Color scheme: C, magenta; N, blue; O, red.

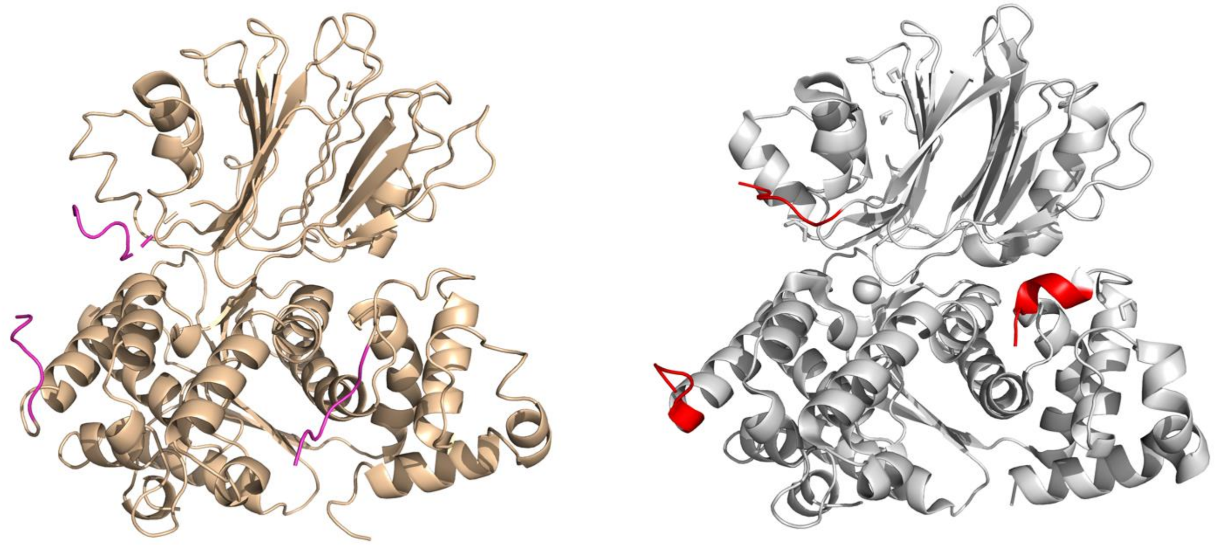

**Figure S9**. **The EM-derived (left) and X-ray crystal (right) structures of the human asparagine synthetase monomer.** Ten residues adjacent to the experimentally undefined regions are colored in magenta and red for the EM-derived and X-ray structures, respectively.

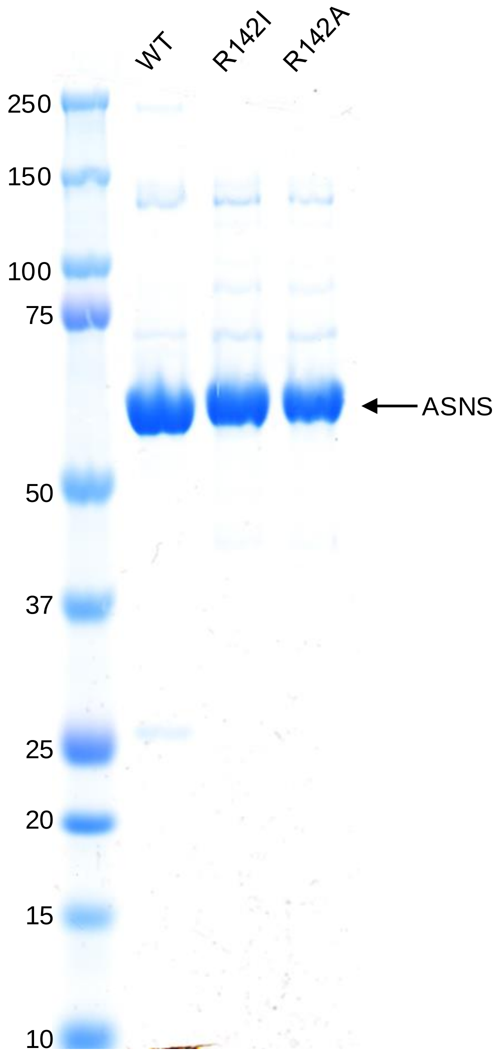

**Figure S10. SDS-PAGE of recombinant WT ASNS, and the recombinant R142I and R142A ASNS variants used in this study.** Approximately 6 µg samples of each recombinant protein were subjected to SDS-PAGE and stained with Coomassie Brilliant Blue. Each ASNS band is indicated by arrow. Left hand column is composed of molecular weight markers (numbers indicate the marker size in kDa). M: molecular weight marker; WT: wild-type ASNS; R142I; R142I variant; R142A; R142A variant.

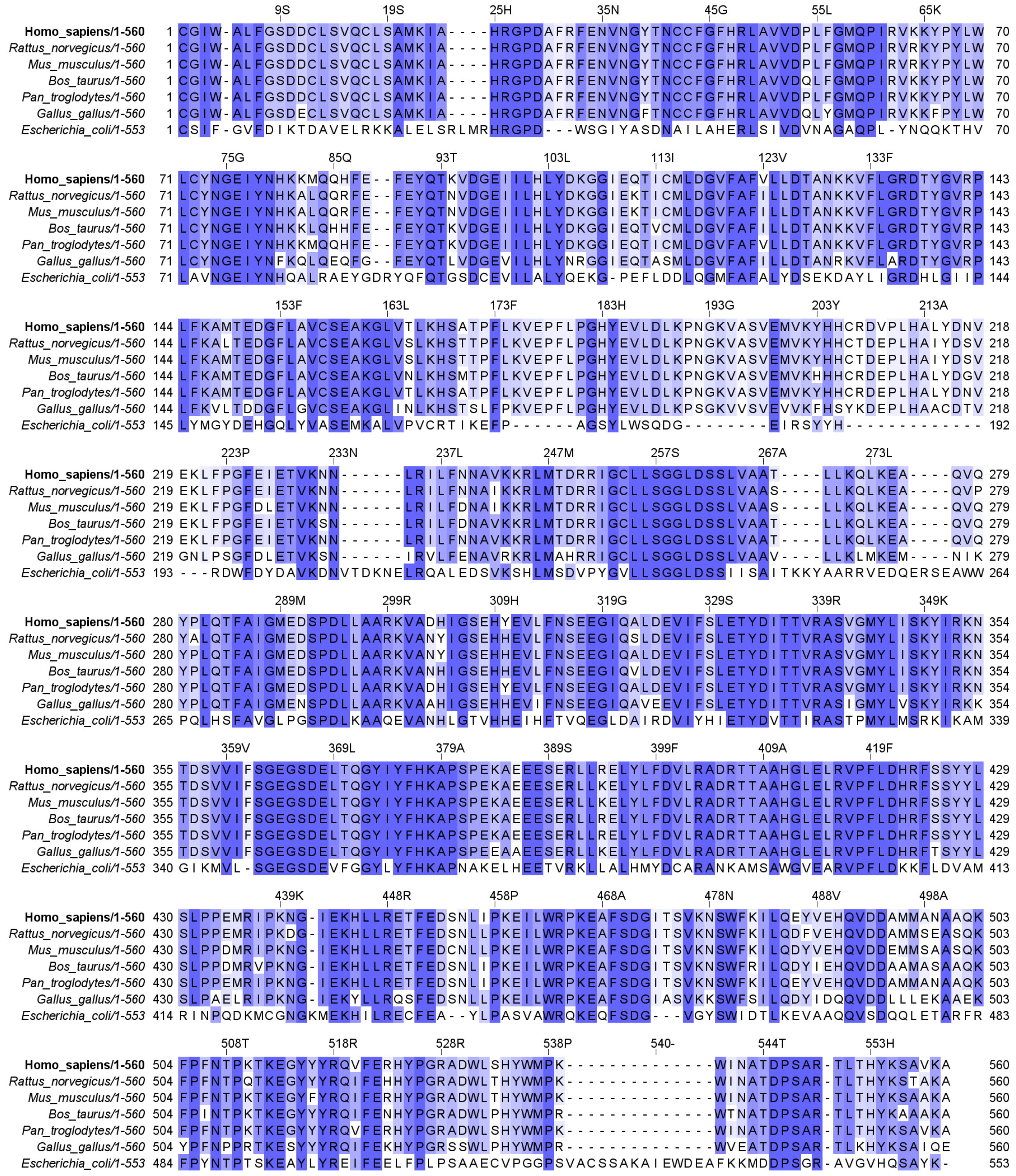

**Figure S11**. **Sequence alignment of the deduced sequences for glutamine-dependent asparagine synthetase in mammals, chickens and *Escherichia coli*.** Shading indicates conserved (dark) and highly conserved (light) residues

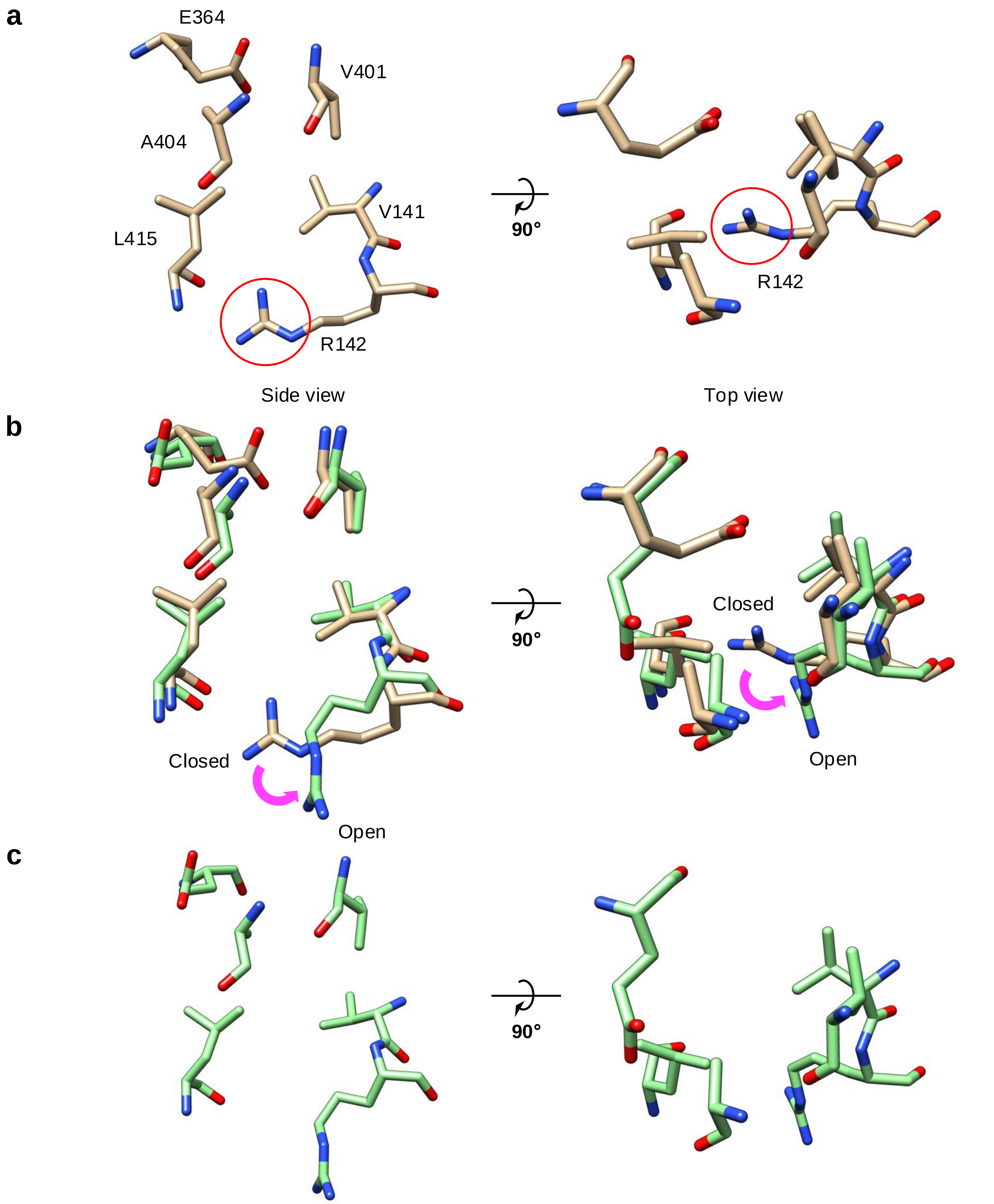

**Figure S12. Example of conformational changes in the ammonia tunnel interconvert the "open and closed" forms in WT ASNS, as detected by 3D variability analysis.**

**a** Critical residues constituting the ammonia tunnel showing how the Arg-142 side-chain (R142) blocks the tunnel in the EM structure

**b** Superimposition of tunnel residues in 3DVA component1 superimposed with those shown in a, indicating how a conformational change of R142 interconverts the open and closed forms of the tunnel

**c** Open conformation detected in the 3DVA component1

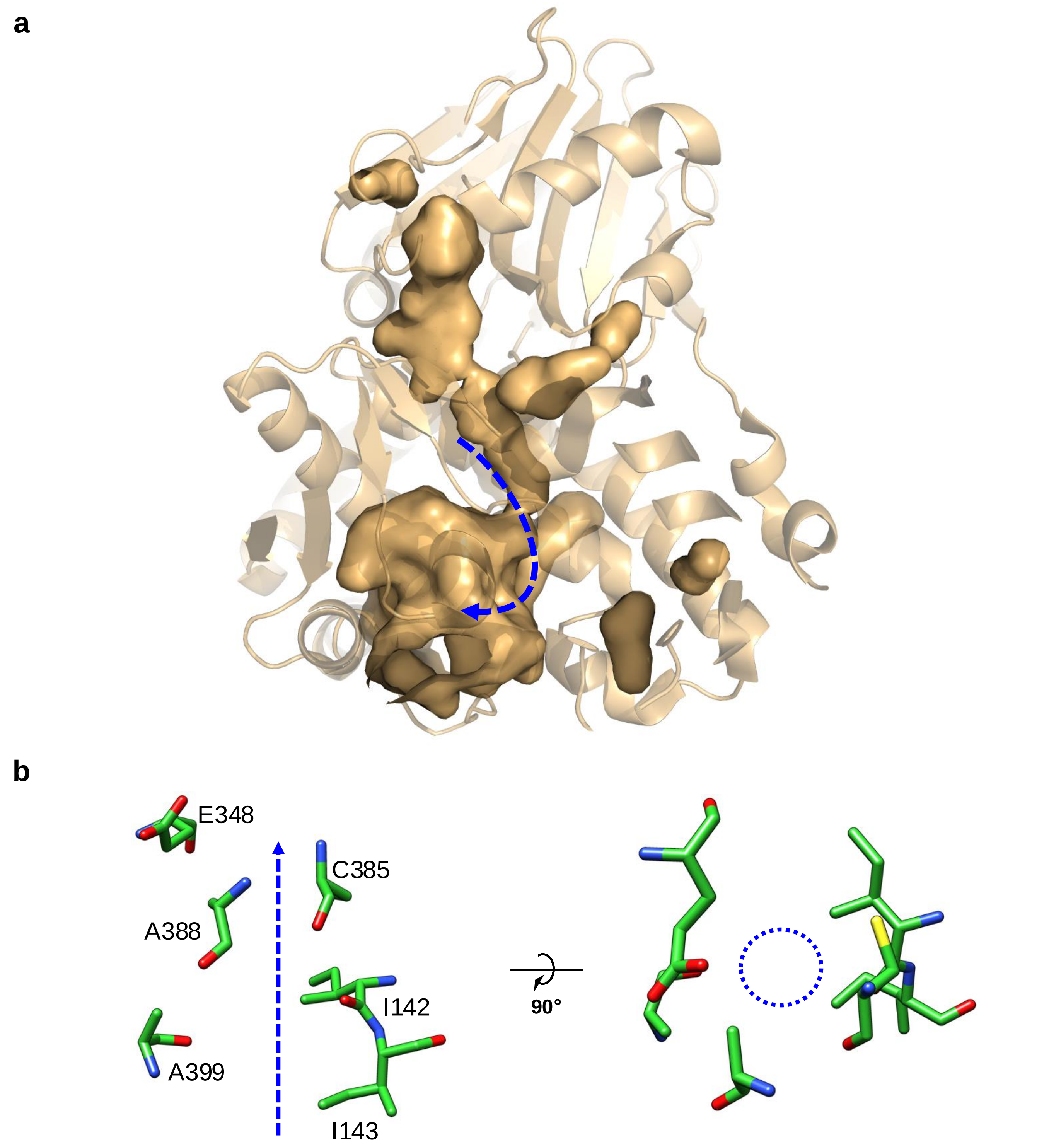

**Figure S13. The open tunnel observed in the X-ray crystal structure of the C1A variant of *Escherichia coli* AS-B complexed with glutamine and AMP. a** The blue dashes represent the location of the open tunnel for $NH_3$ translocation from the N-terminal glutaminase site to the C-terminal synthetase site. **b** Critical residues constituting the tunnel: side view (left) and top view (right). The dashed blue arrow and circle illustrate an open tunnel configuration.

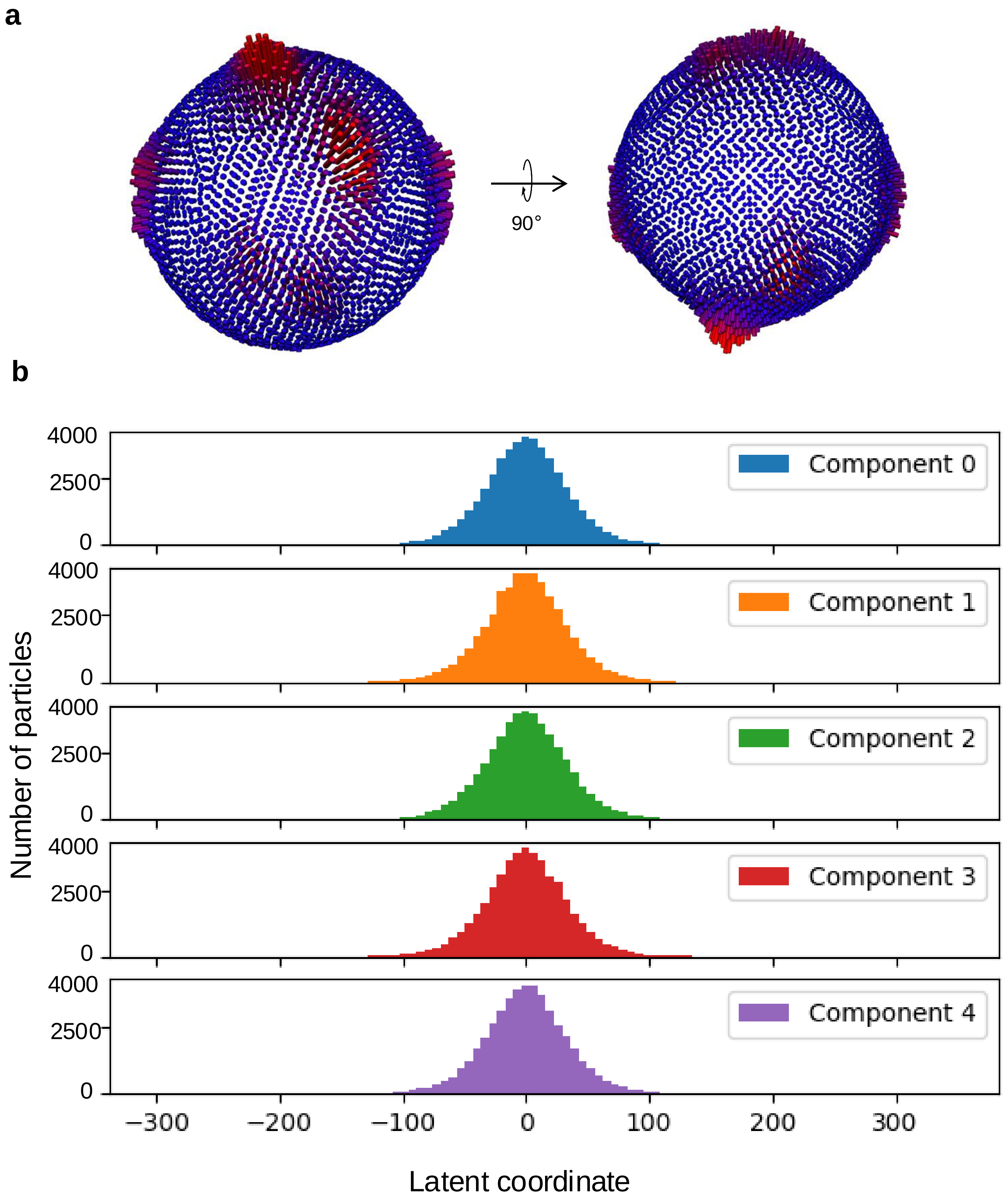

**Figure S14. PCA and particle distributions.**
**a** 3D representation of particle distribution used in the reconstruction; 2 views at 90° are represented. Red cylinders indicate the amount of particle in each rotation angle for the reconstruction.
**b** Particle distribution in latent space, represented per principal component as a function of latent coordinate.

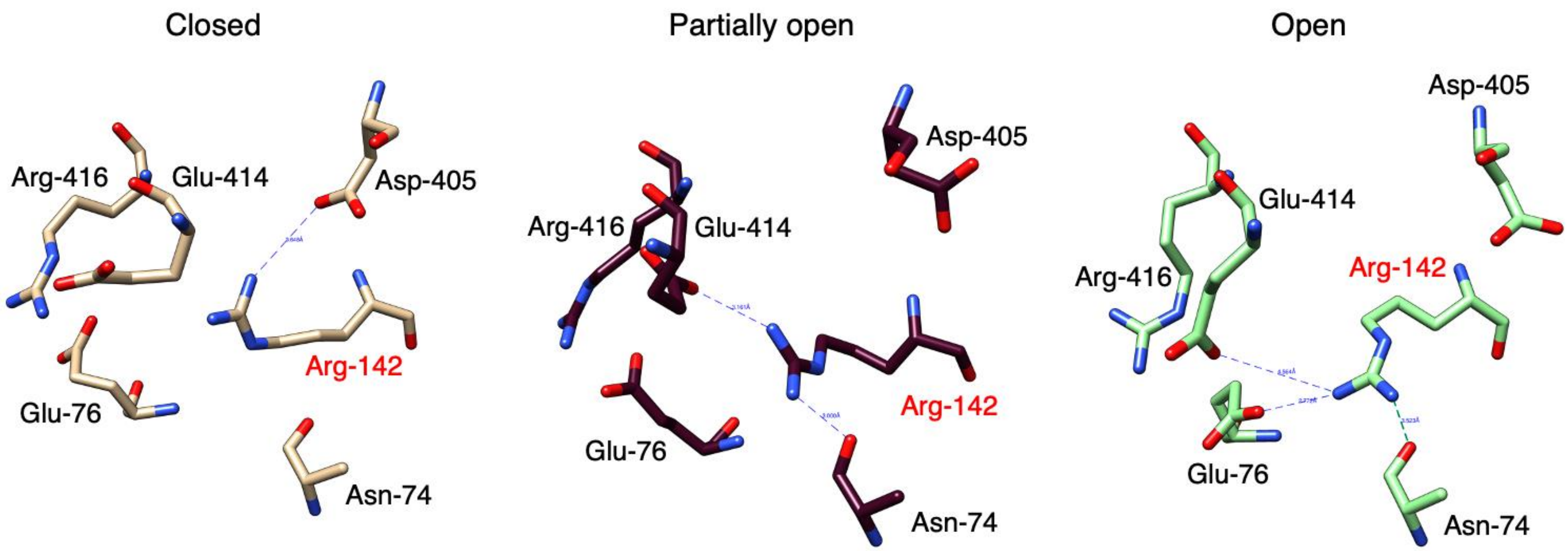

**Figure S15**. **Distance-based definitions of closed, open, and partially open Arg-142 conformations seen in the 3DVA analysis.** (Left) In the closed conformation, blocking ammonia access to the tunnel, Arg-142 only makes a direct electrostatic interaction with the carboxylate of Asp-405. (Middle) In the partially open conformation, the Arg-142/Asp-405 interaction is disrupted, and the side chains move apart. As a result, the guanidinium group becomes closer to the side chains of Asn-74 and Glu-414. (Right) In the fully open conformation, which permits ammonia access to the tunnel, the guanidinium moiety of Arg-142 forms an additional interaction with the carboxylate of Glu-76. The Arg-142/Asp-405 and Arg-142/Glu-414 distances can therefore be used as a measure of whether the side chain is in the closed conformation (shorter/longer, respectively) or open conformation (shorter/longer, respectively).

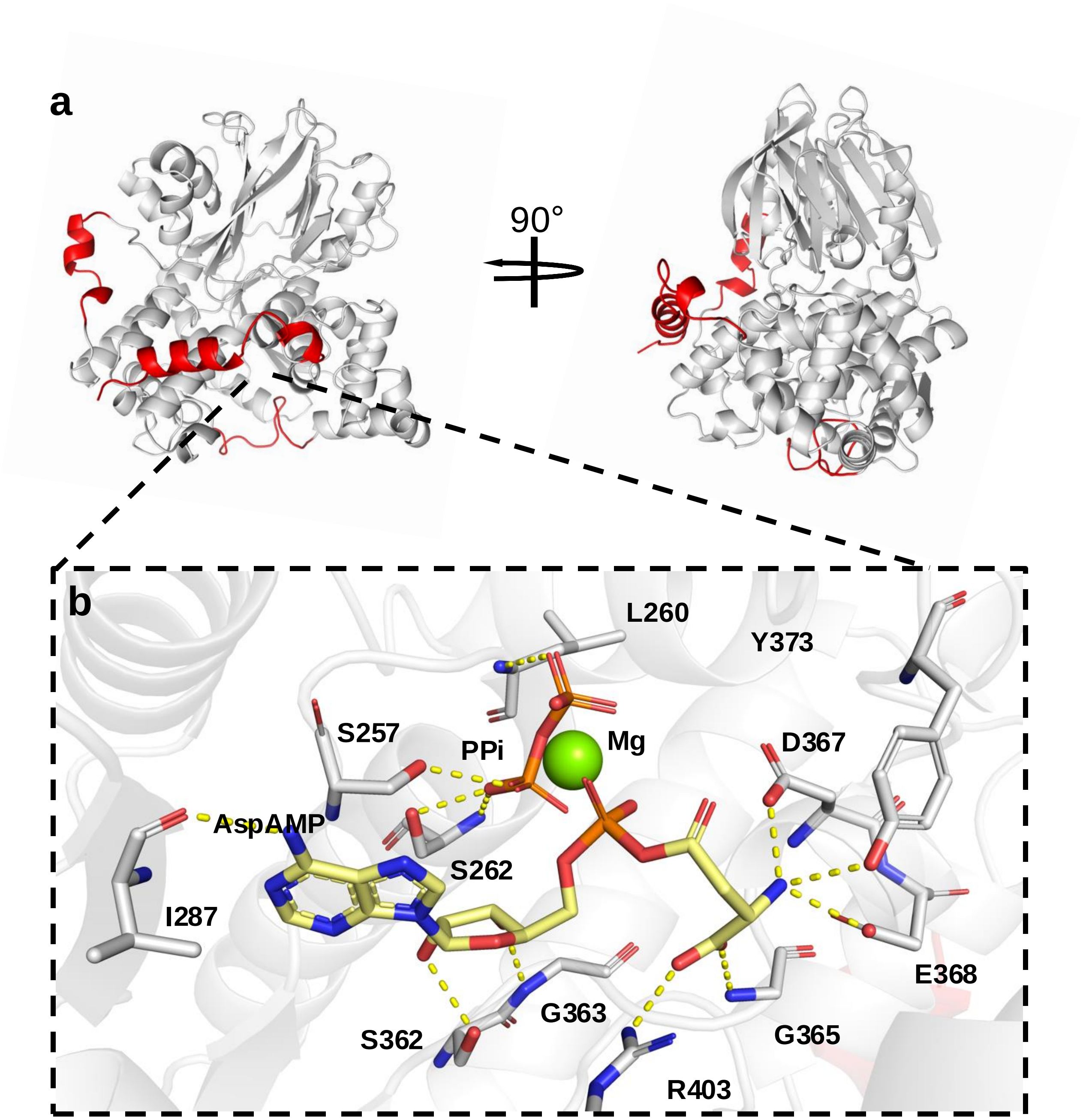

**Figure S16**. **Computational models constructed from the Robetta structure of human ASNS. a** Cartoon representation of the full-length model of human apo-ASNS used in the computational studies, showing the reconstructed loops (red). **b** Close-up of the synthetase active site in the computational model of the human ASNS/β-aspartyl-AMP/$PP_i$ ternary complex. Key protein residues (C, grey) and the ligands (C, yellow) are rendered as sticks and $Mg^{2+}$ is shown by the green sphere. Intermolecular hydrogen bonds are indicated by the dashed yellow lines. (Note that these models were also used to generate initial structures for the apo-R142I variant and its β-aspartyl-AMP/$PP_i$ ternary complex).

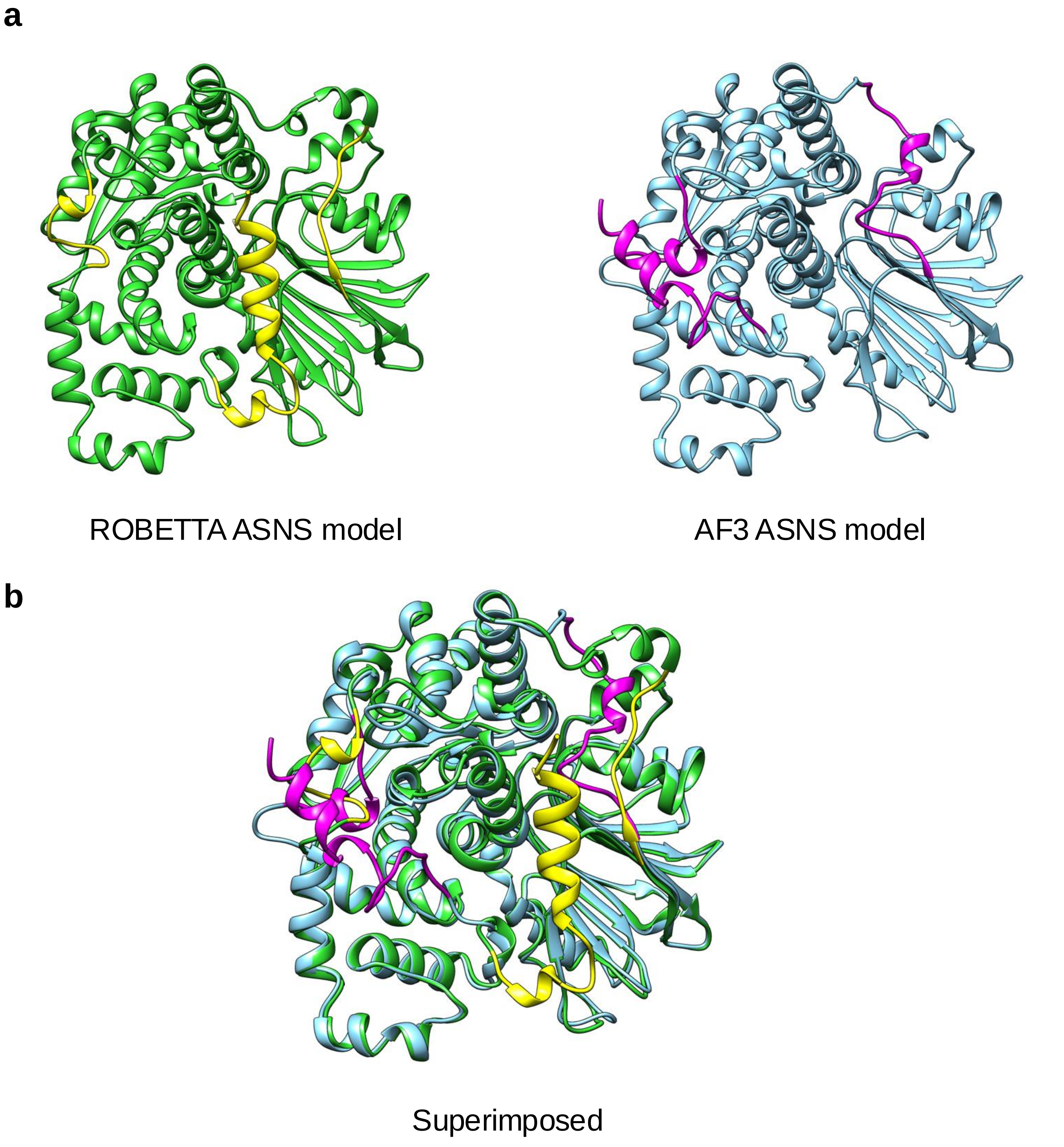

**Figure S17. Comparison of ROBETTA-generated and Alphafold3 (AF3)-generated models for the WT human apo-ASNS monomer. a** Ribbon representation of the ROBETTA model (lime green), and the AF3-generated model (sky blue). Loop regions (residues 201-220, 465-475) as well as the C-terminal tail (residues 536-560), which are not seen in the X-ray or cryo-EM structures are colored in yellow for the ROBETTA model and magenta for the AF3 model. **b** Superimposed models of ROBETTA-generated and AF3-generated models (RMSD: 0.629 Å). The major difference is associated with the position of the C-terminal region in the two models.

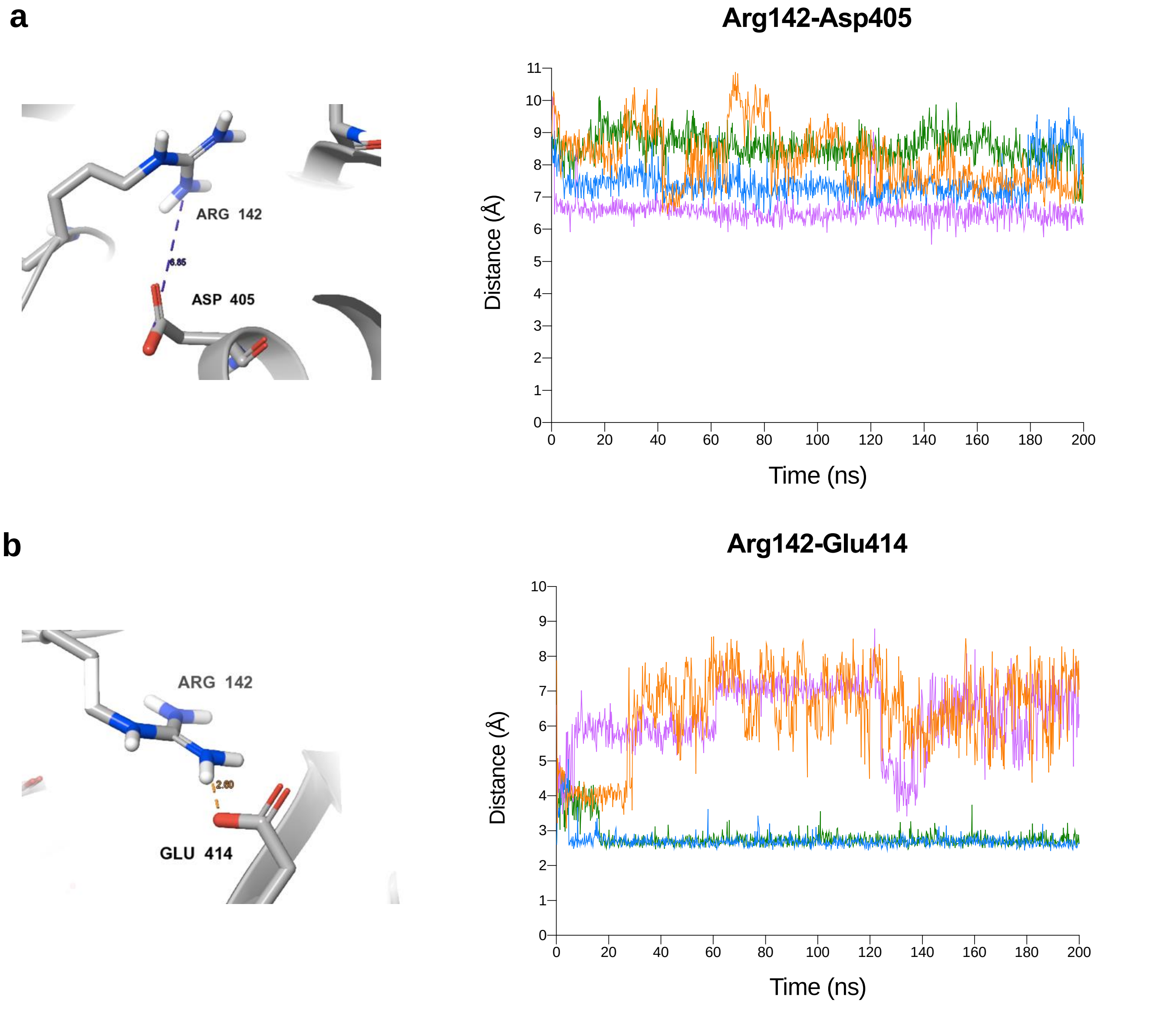

**Figure S18**. **Replicate 200 ns MD simulations of the apo-ASNS monomer. a** Plots of the distance between Arg-142 and Asp-405 (shown on the left) in the four replicates of the 200 ns MD simulations of the apo-ASNS monomer. **b** Plots of the distance between Arg-142 and Glu-414 (shown on the left) in the four replicates of the 200 ns MD simulations of the apo-ASNS monomer. Distances are calculated between the NH2 nitrogen atoms of Arg-142 and carboxylate oxygen atoms of Asp-405 or of Glu-414, respectively, as indicated by the dashed lines. Shorter Arg-142/Asp-405 (longer Arg-142/Glu-414) distances correspond to Arg-142 adopting the “closed” conformation in the cryo-EM structure that blocks ammonia access. Similarly, longer Arg-142/Asp-405 (shorter Arg-142/Glu-414) distances correspond to Arg-142 adopting the “open” conformation in the cryo-EM structure that permits ammonia access.

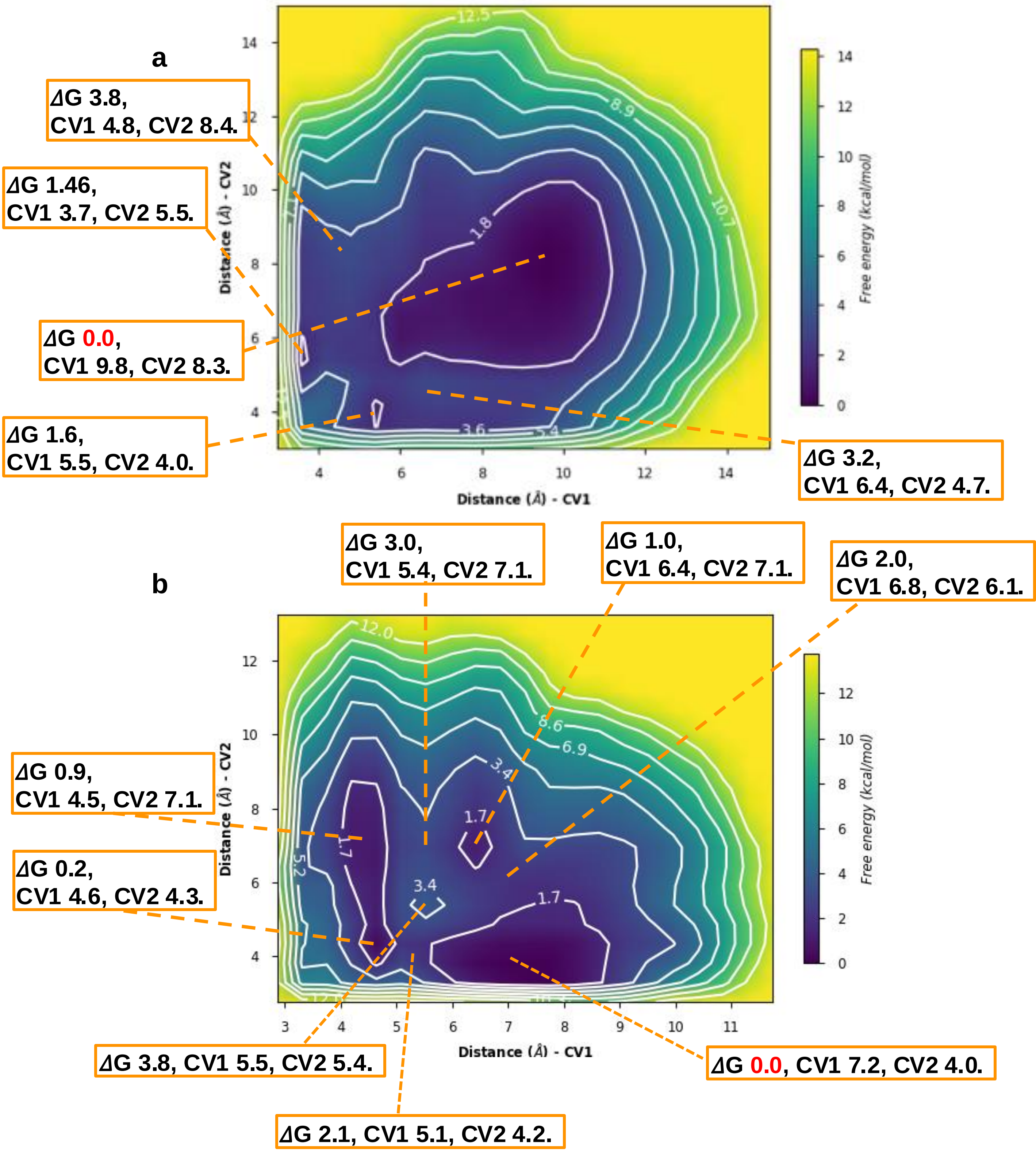

**Figure S19. Metadynamics-based free energy surfaces for the Arg-142 side chain in (a) apo-ASNS and (b) the WT ASNS/β-aspartyl-AMP/$PP_i$ ternary complex.** Two distance-based collective variables were defined for these simulations (Supplementary Figs. S15), which correspond to the distances between the center-of-mass (COM) of the guanidinium group in Arg-142 and the COM of the carboxylate moieties in either Asp-405 (CV1) or Glu-414 (CV2).

**a**

**b**

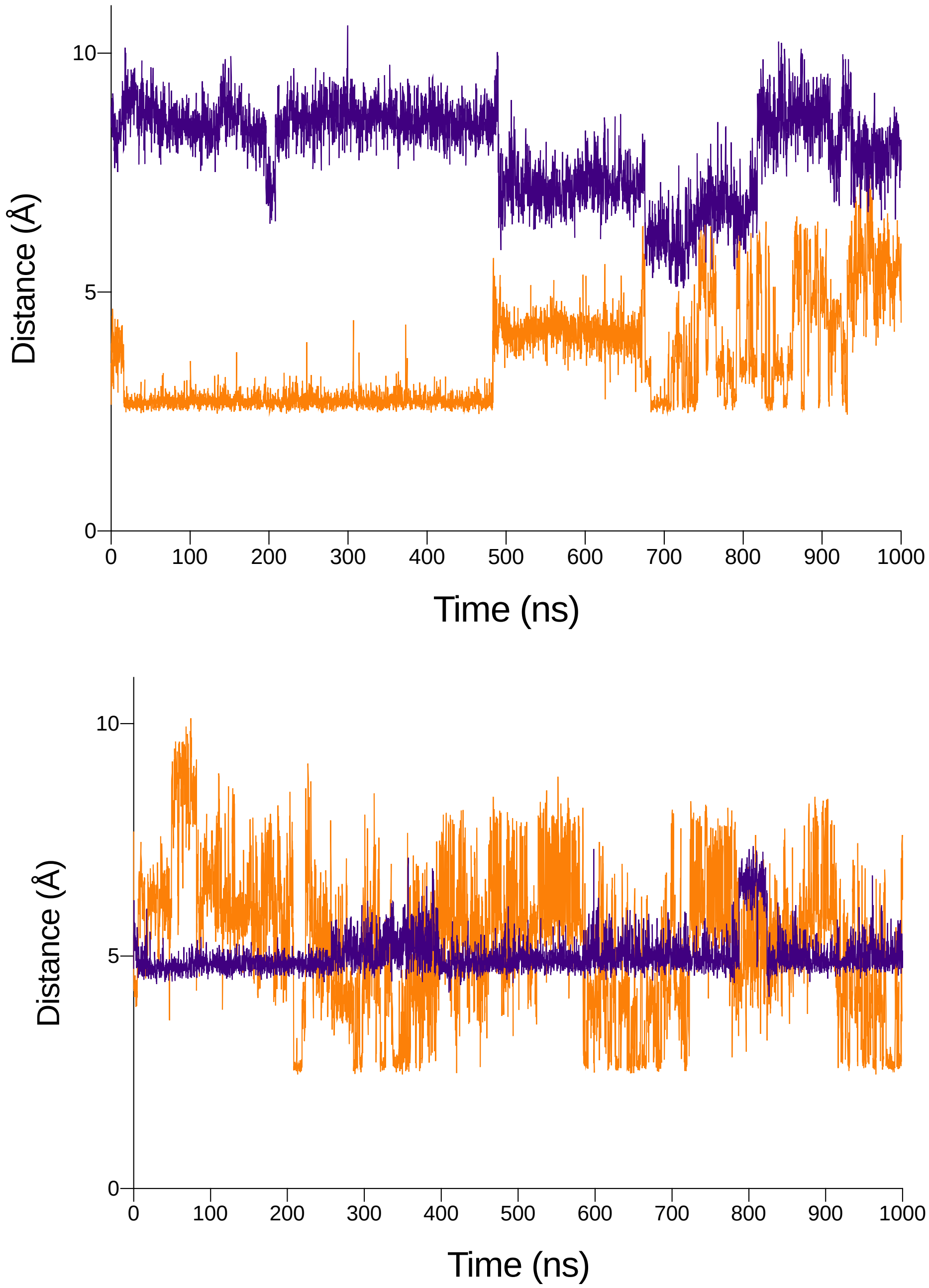

**Figure S20**. **Microsecond simulations of apo-ASNS and the WT ASNS/β-aspartyl-AMP/$PP_i$ ternary complex in their monomeric form. a** Plots of the distance between Arg-142 and Glu-414 (orange) and between Arg-142 and Asp-405 (purple) in the microsecond MD simulation of the full-length apo-ASNS monomer. **b** Plots of the distance between Arg-142 and Glu-414 (orange) and between Arg-142 and Asp-405 (purple) in the microsecond MD simulation of the WT ASNS/β-aspartyl-AMP/$PP_i$ ternary complex. All distances are defined as described in the legend to Fig. S17.

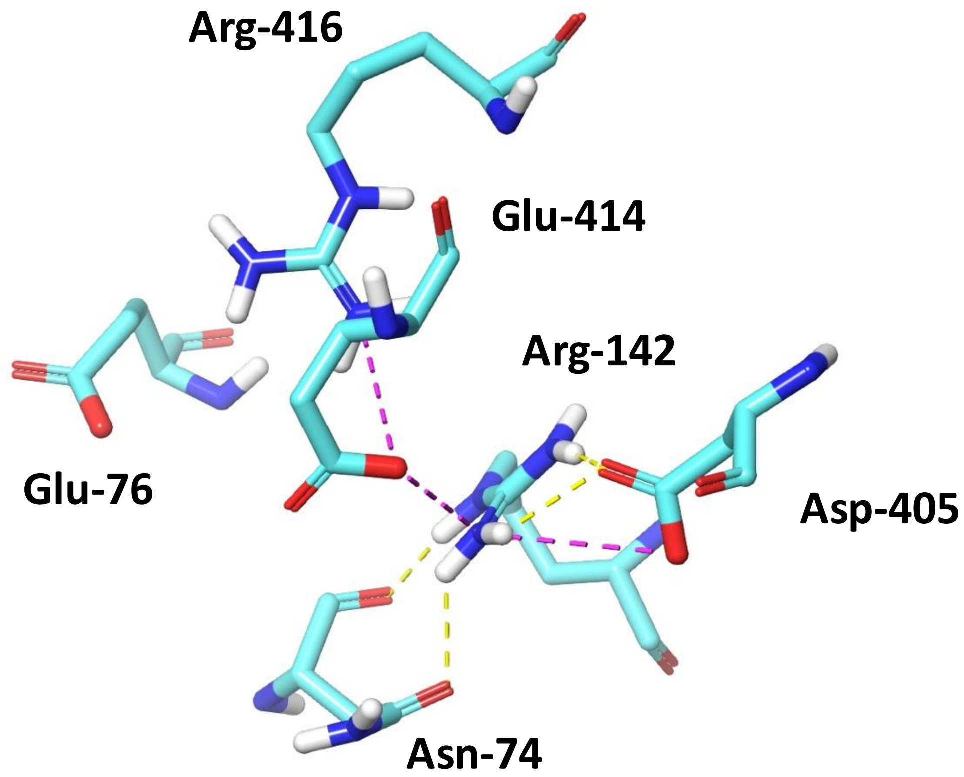

**Figure S21**. **Residue network showing the Arg-142/Asn-74 interaction.** Structure is taken from a snapshot (t = 200 ns) in the MD trajectory of apo-ASNS. Hydrogen bonds and salt bridges are indicated by yellow and violet dashed lines, respectively. Color scheme: C, cyan; H, white; N, blue; O, red.

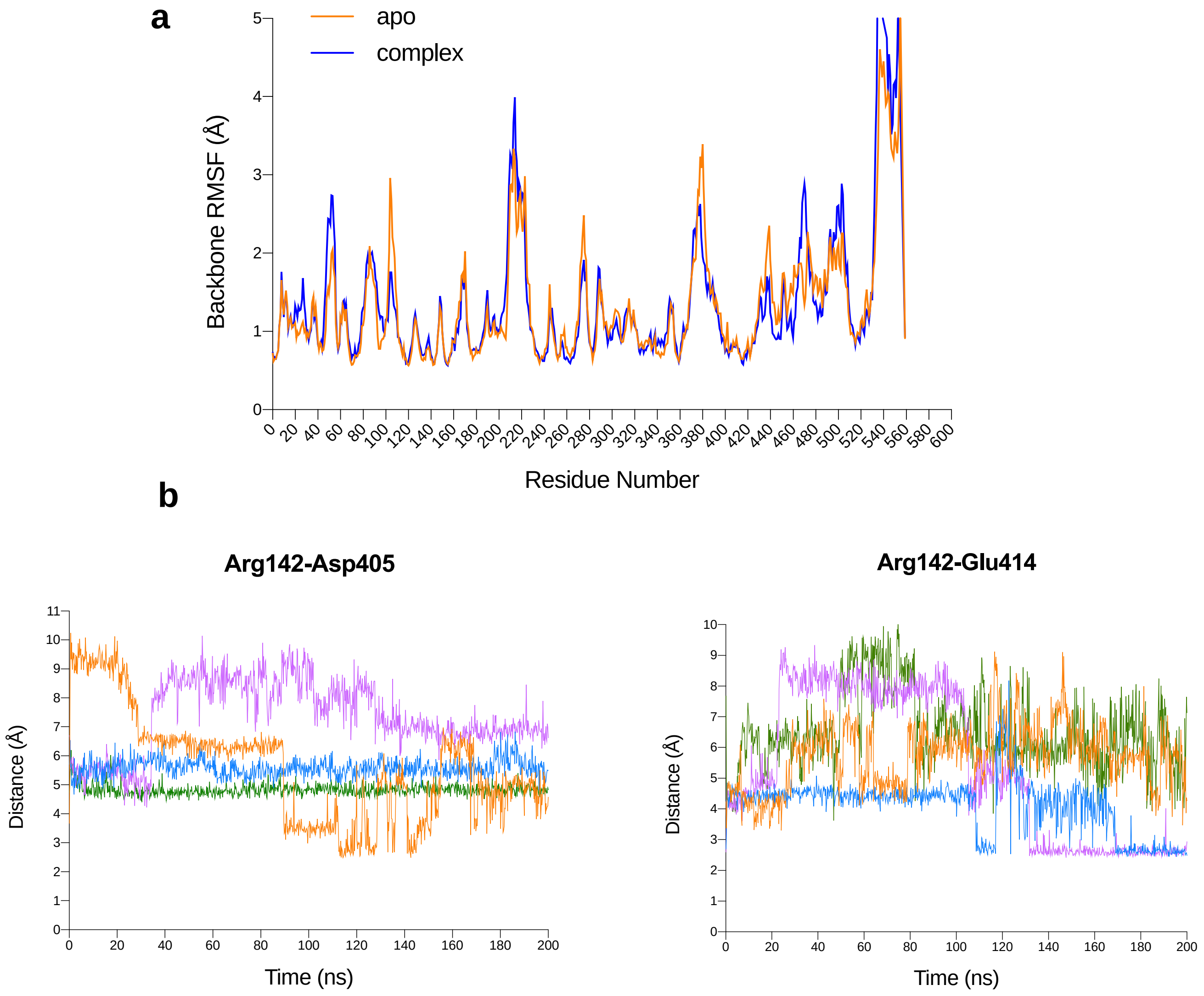

**Figure S22. 200 ns MD simulations of the WT ASNS/β-aspartyl-AMP/PP$_i$ ternary complex**. **a** Comparison of backbone RMSF values computed for the apo-ASNS monomer (orange) and the WT ASNS/β-aspartyl-AMP/PP$_i$ ternary complex (blue). **b** Replicate 200 ns MD simulations of the WT ASNS/β-aspartyl-AMP/PP$_i$ ternary complex. (left) Plots of the distance between Arg-142 and Asp-405 and (right) between Arg-142 and Glu-414 in the four replicates. All distances are defined as described in the legend to Fig. S17.

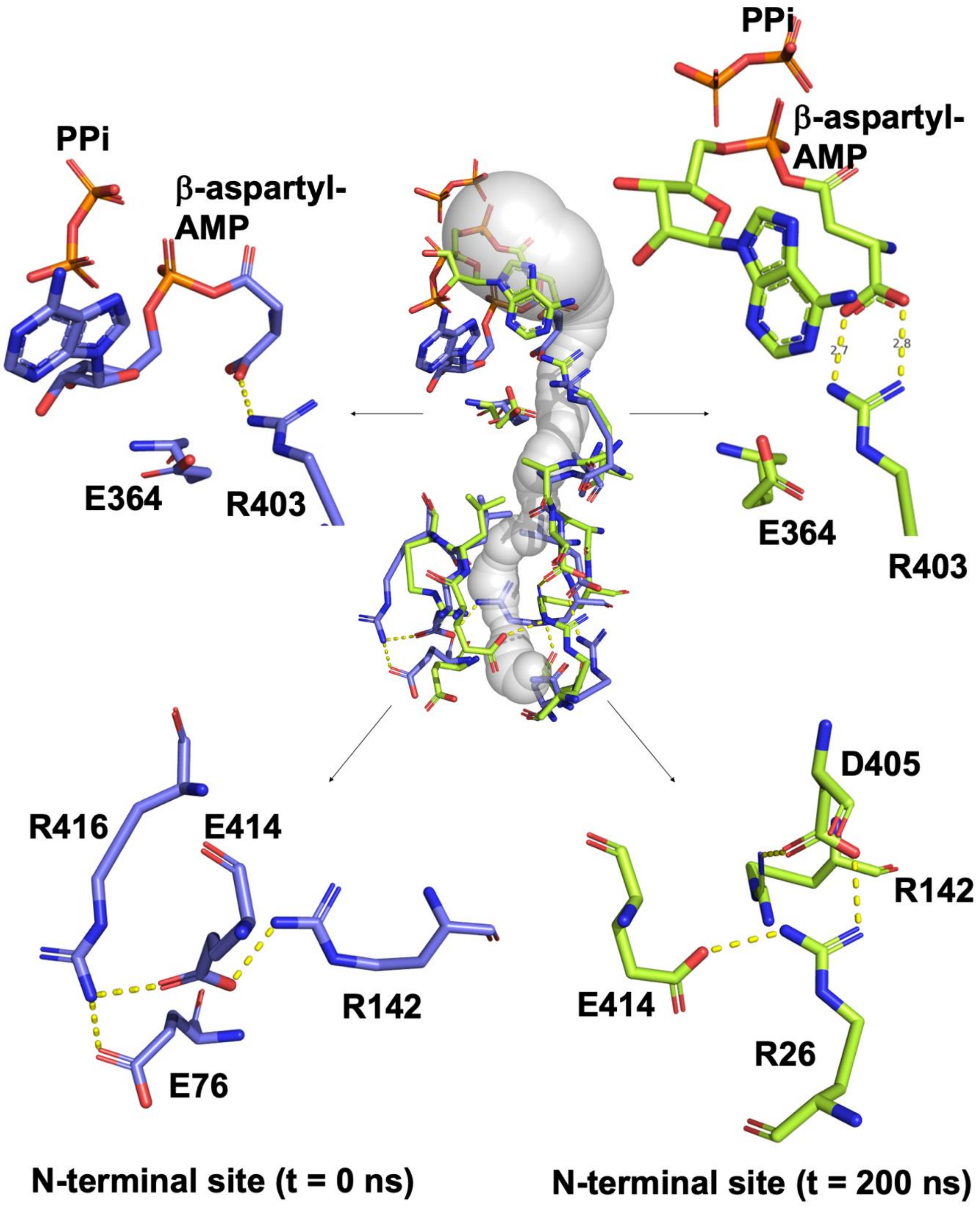

**Figure S23**. **Communication network between the N-terminal active site and the C-terminal active site as seen in the MD trajectory of the WT ASNS/β-aspartyl-AMP/PP$_i$ ternary complex.** Residues are shown in sticks; carbon atoms are shown in purple and lime for the t = 0 ns and t = 200 ns snapshots, respectively. Oxygen atoms are shown in red, and nitrogen atoms are shown in blue. Yellow dashed lines show residue interactions within 2.8 Å. The tunnel present in the final snapshot of the 200 ns simulation is shown in gray.

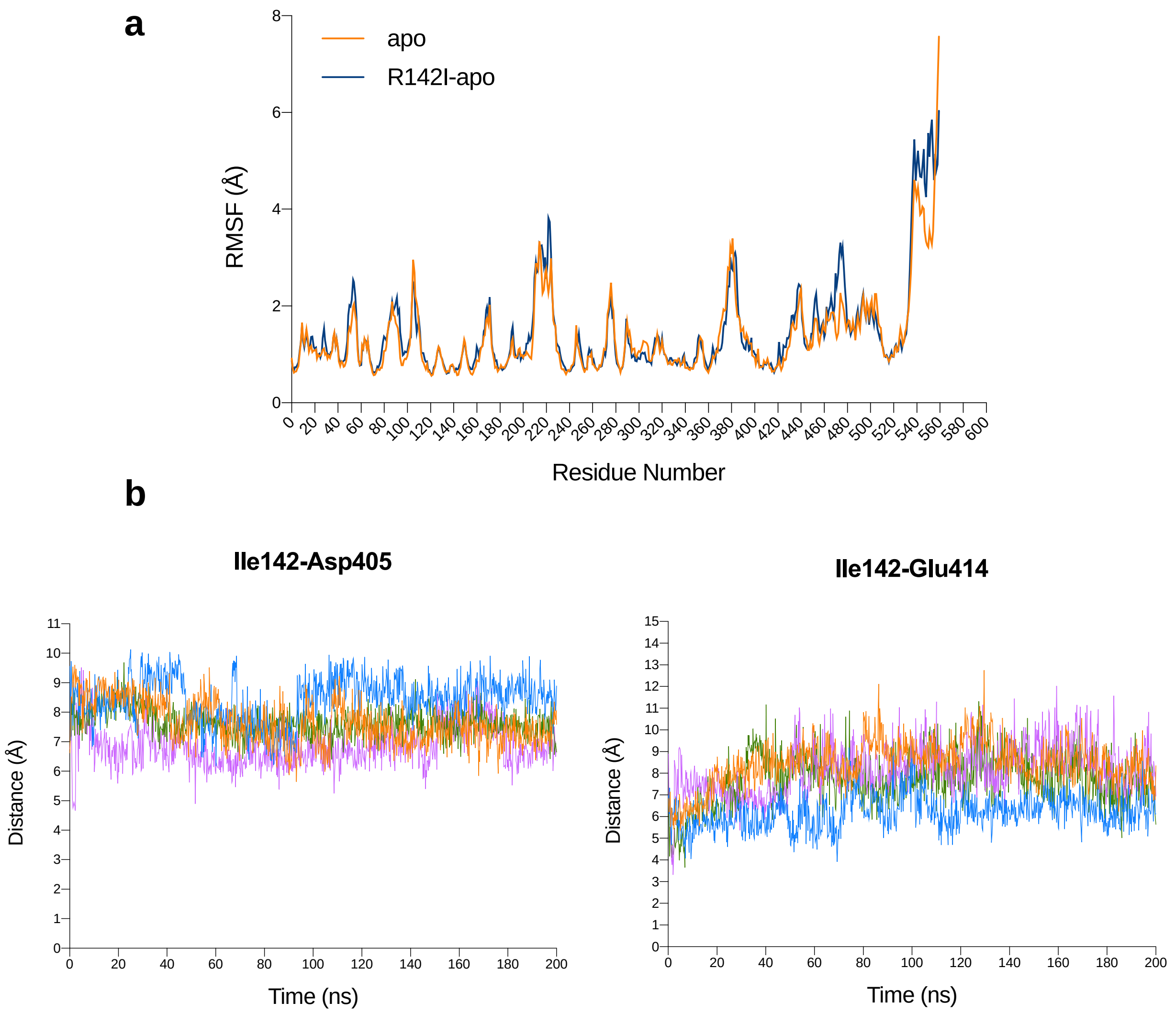

**Figure S24**. **200 ns MD simulations of the apo-R142I variant. a** Comparison of backbone RMSF values computed for the apo-ASNS monomer (orange) and the apo-R142I variant (blue). **b** Replicate 200 ns MD simulations of the apo-R142I variant. (left) Plots of the distance between Ile-142 and Asp-405 and (right) between Ile-142 and Glu-414 in the four MD simulation replicates. Distances were calculated between the delta carbon atom of Ile-142 and one of the carboxylate oxygen atoms of Asp-405 or of Glu-414, respectively. See Fig. S17 for the location of these oxygen atoms.

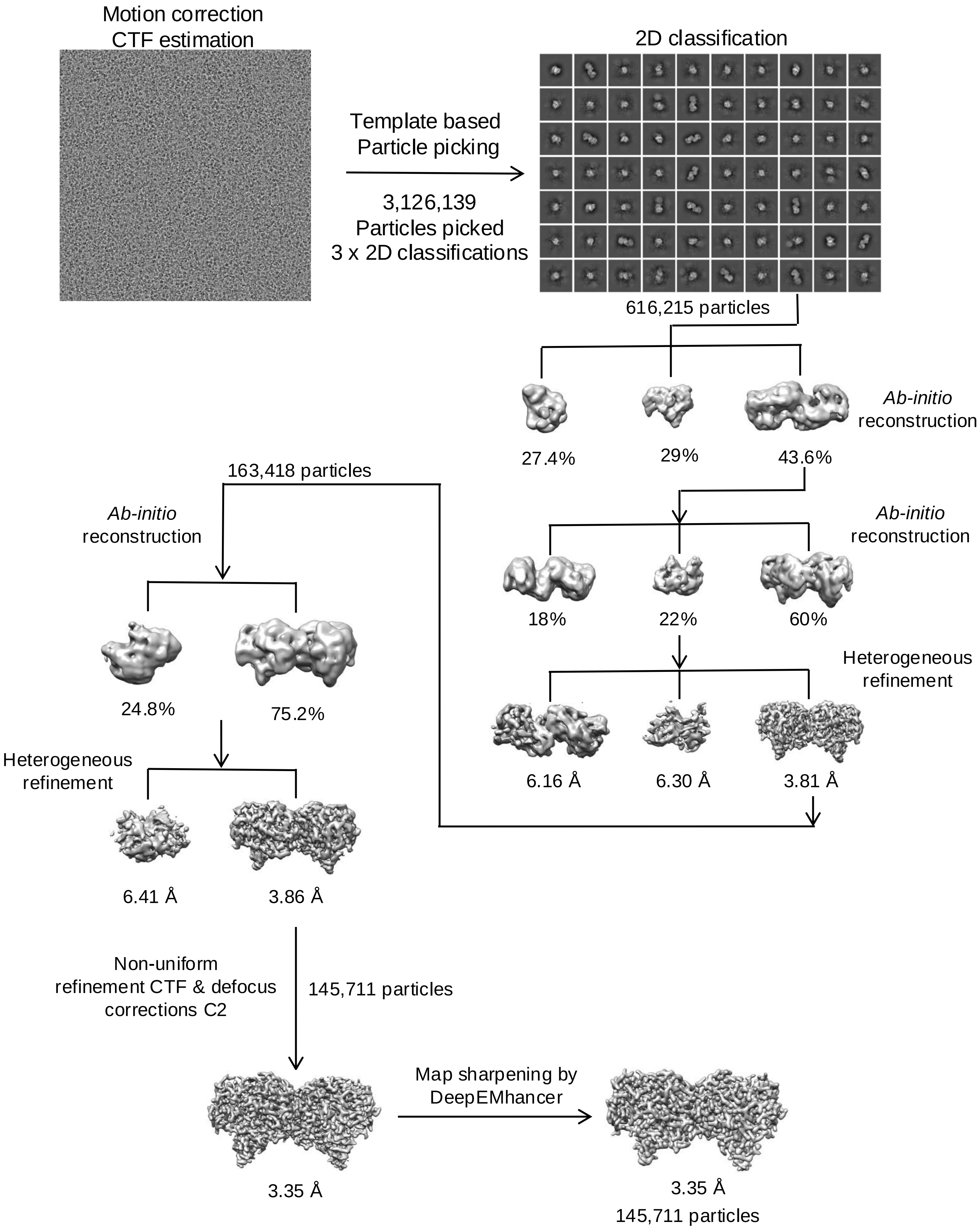

**Figure S25. Summary of the cryo-EM data processing workflow of ASNS R142I variant using cryoSPARC v4.4.1.**

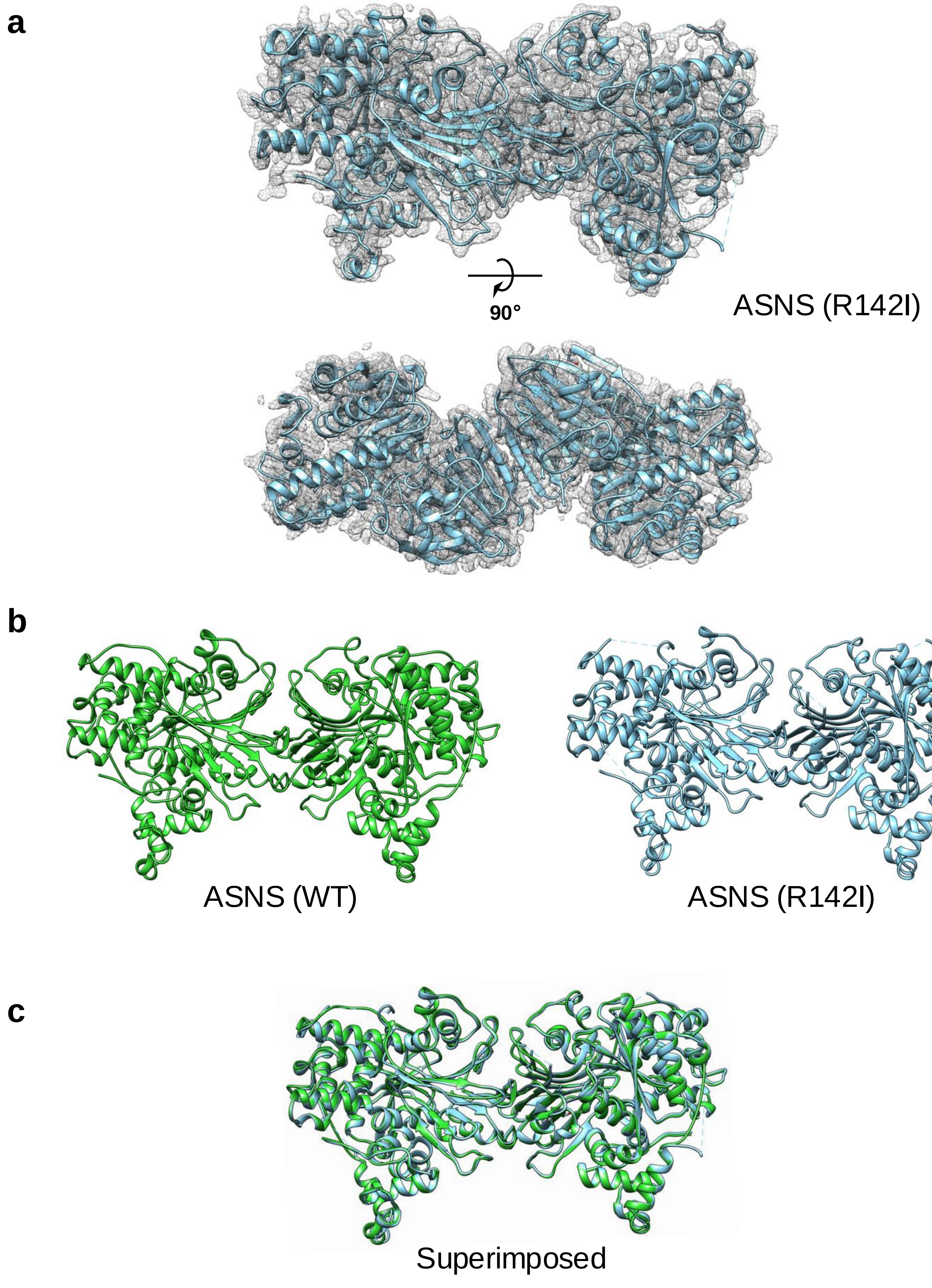

**Figure S26. Structure of the R142 ASNS variant and its comparison to ASNS WT**
**a** Ribbon representation of the structure of the R142I ASNS variant (PDB: 9B6C) colored in sky blue with its EM map (EMD-44253) in grey. **b** Comparison between WT vs. R142I variant structures: side view of the WT model (PDB: 8SUE) colored in lime green on the left, and the R142I variant model (PDB: 9B6C) in sky blue on the right. **c** Superimposed models of WT and the R142I variant (RMSD: 0.483 Å)

### a Gold standard FSC curve

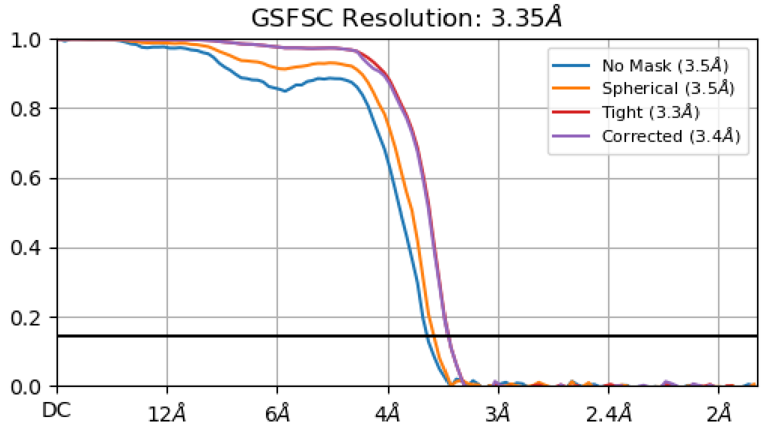

### b Viewing distribution plots

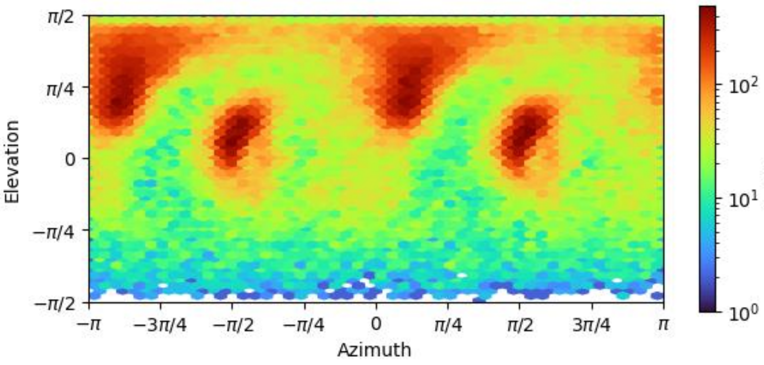

### c 3D FSC

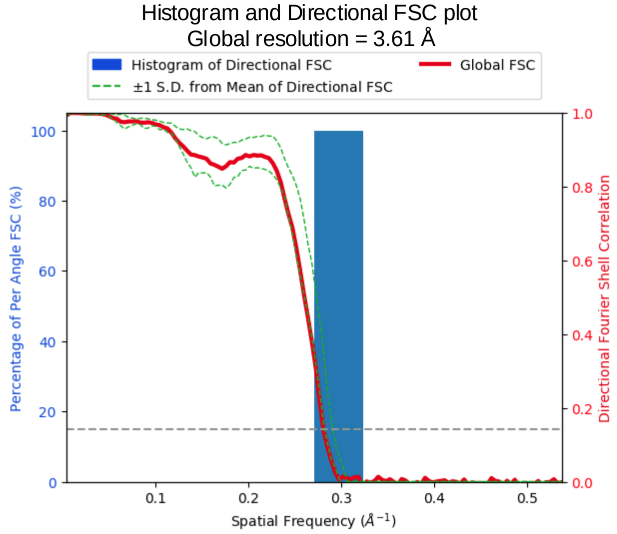

### d Local resolution

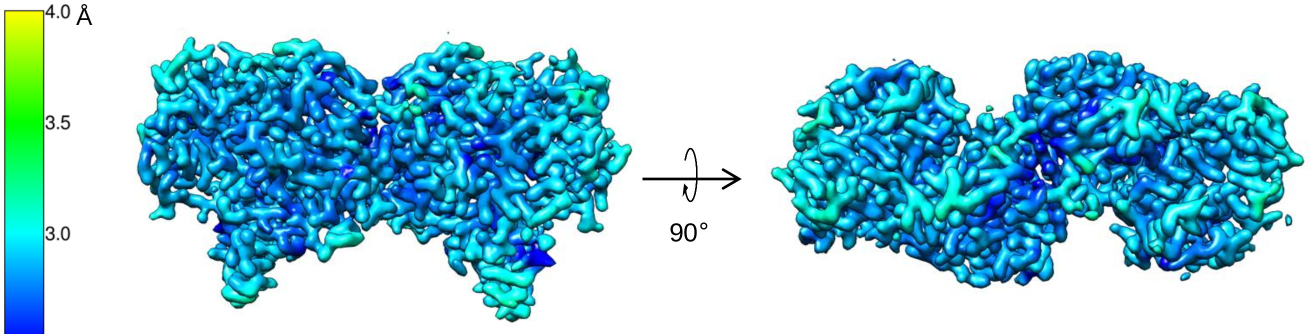

**Figure S27. Assessment of the quality of cryo-EM map of R142I ASNS variant.**
**a** Gold standard FSC curve for the map of R142I variant generated in cryoSPARC v4.4.1.
**b** Viewing distribution plots for the map generated in cryoSPARC v4.4.1.
**c** 3D FSC: histogram and directional FSC plot for map global resolution are shown.
**d** Local resolution of the map (color scale shown on the left).

### a Representative densities and corresponding models

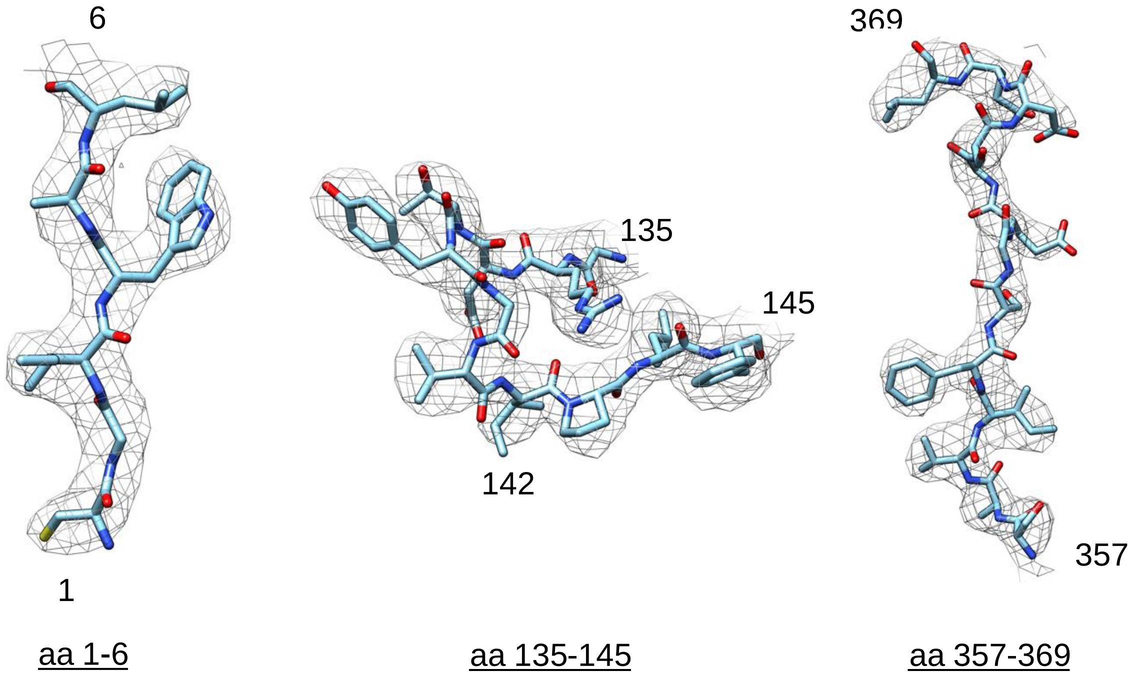

### b Map-model FSC curve

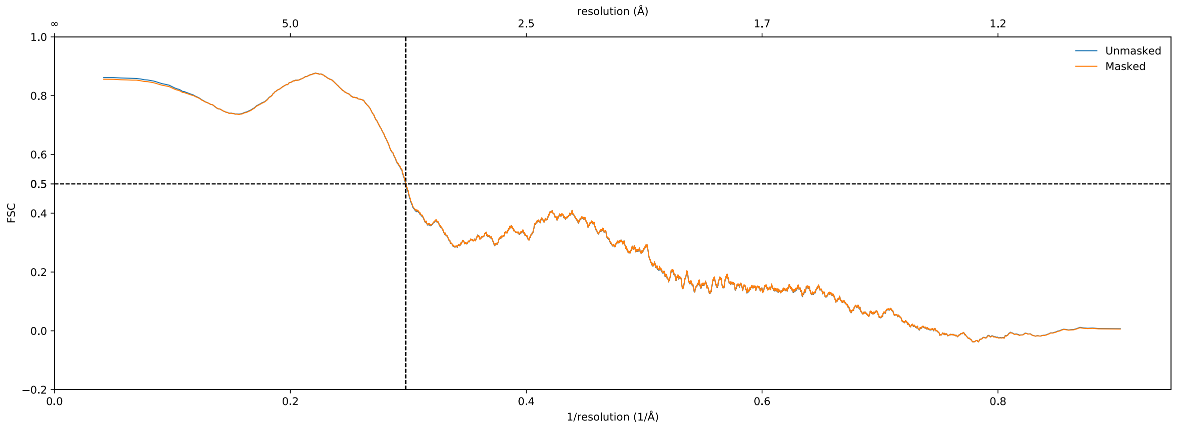

**Figure S28. Validating the cryo-EM structure of ASNS R142I variant : map and its model.**
**a** Representative density of the map for residues in the N-terminal catalytic region (aa 1-6), the ammonia tunnel region (aa 135-145), and the C-terminal active site (aa 357-369).
**b** Model-to-map FSC curves generated in Phenix.

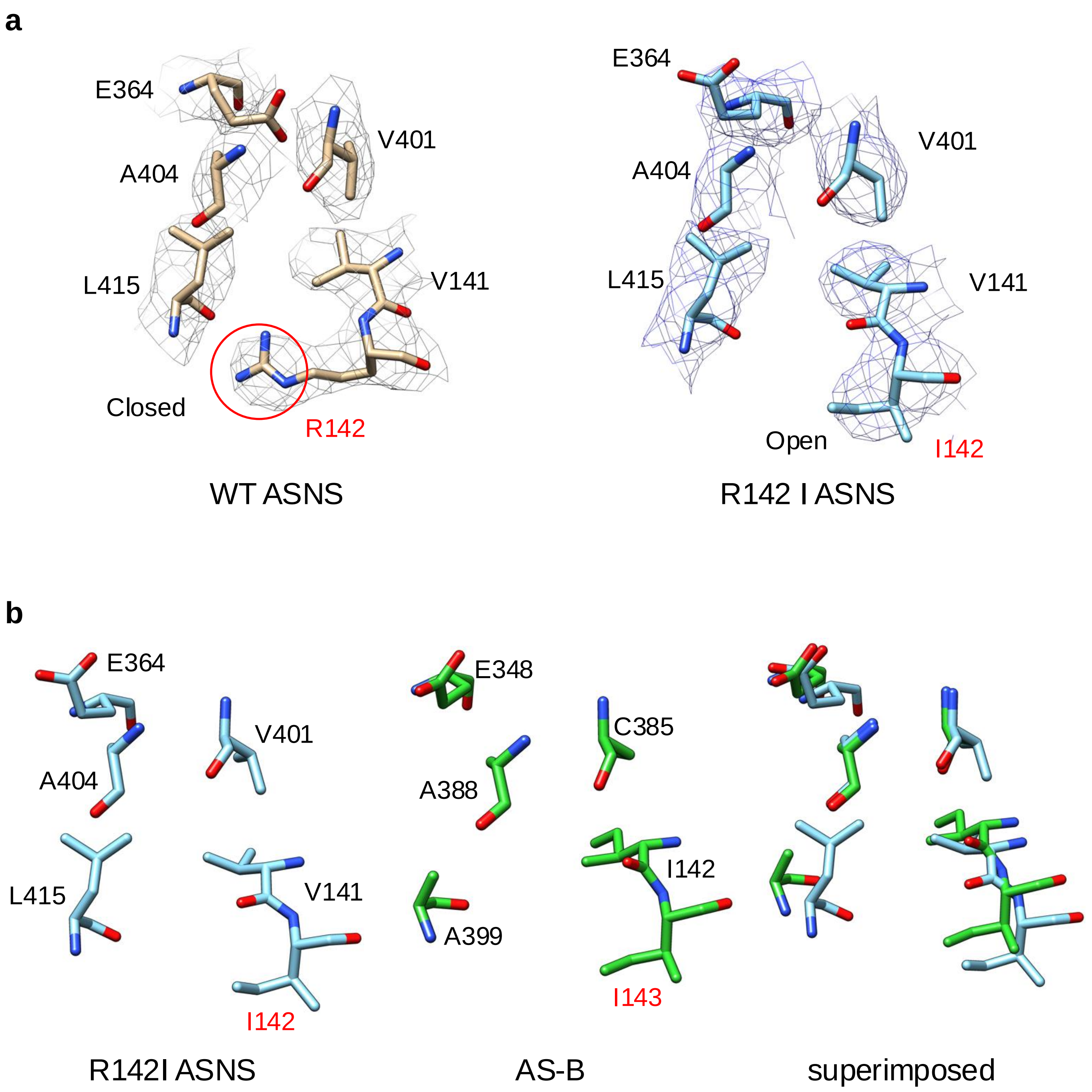

**Figure S29. Structural comparison in tunnel region between the R142I variant and AS-B.**
**a** Comparison of tunnel residues between WT ASNS (left) and the R142I variant (right).
**b** Comparison of tunnel residues between the R142I variant (left) and AS-B (middle), superimposed (right). R142I variant residues, V141, I142, E364, V401, A404, L415 correspond to the residues of bacterial enzyme AS-B, I142, I143, E352, C385, A388, and A399. Residues of WT ASNS are colored in brown, the R142I variant in sky blue, and AS-B in lime green.

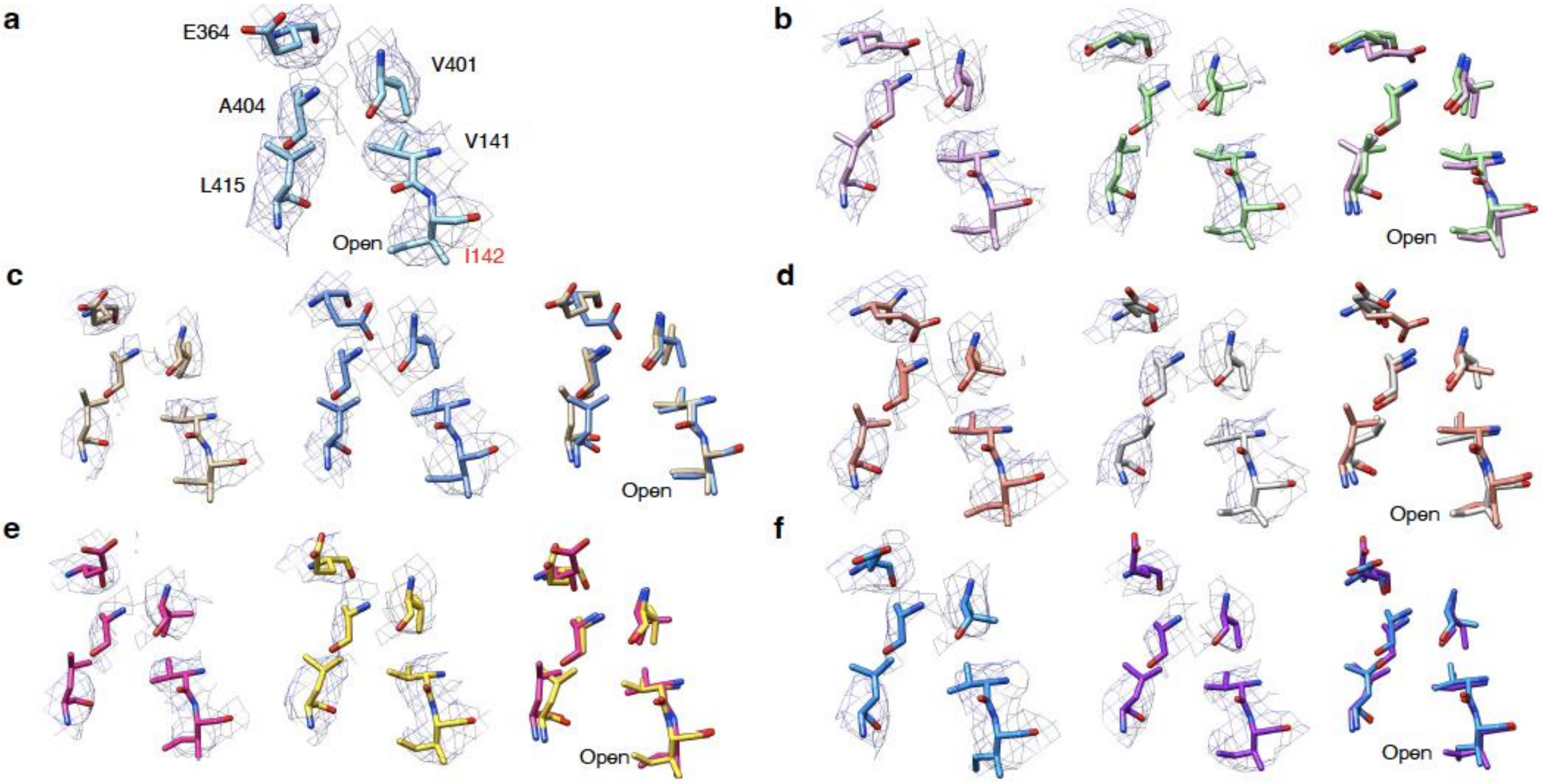

**Figure S30. PCA-derived structures of the ammonia tunnel in the human R142I variant obtained from 3D variability analysis and 3D refinement. a** Side view of critical residues (Val-141, Ile-142, Glu-364, Val-401, Ala-404, Val-414, Leu-415) constituting the ammonia tunnel derived from the consensus EM map. Mutated residue Ile-142 is colored in red. Side view of critical residues constituting ammonia tunnel derived from the variable EM maps derived from 3DVA. Critical residues constituting ammonia tunnel derived from variable maps at frame 1 (left, one end) and frame 20 (middle, the other end) are superimposed (right): **b** PCA component 0, **c** PCA component 1, **d** PCA component 2, **e** PCA component 3, **f** PCA component 4.

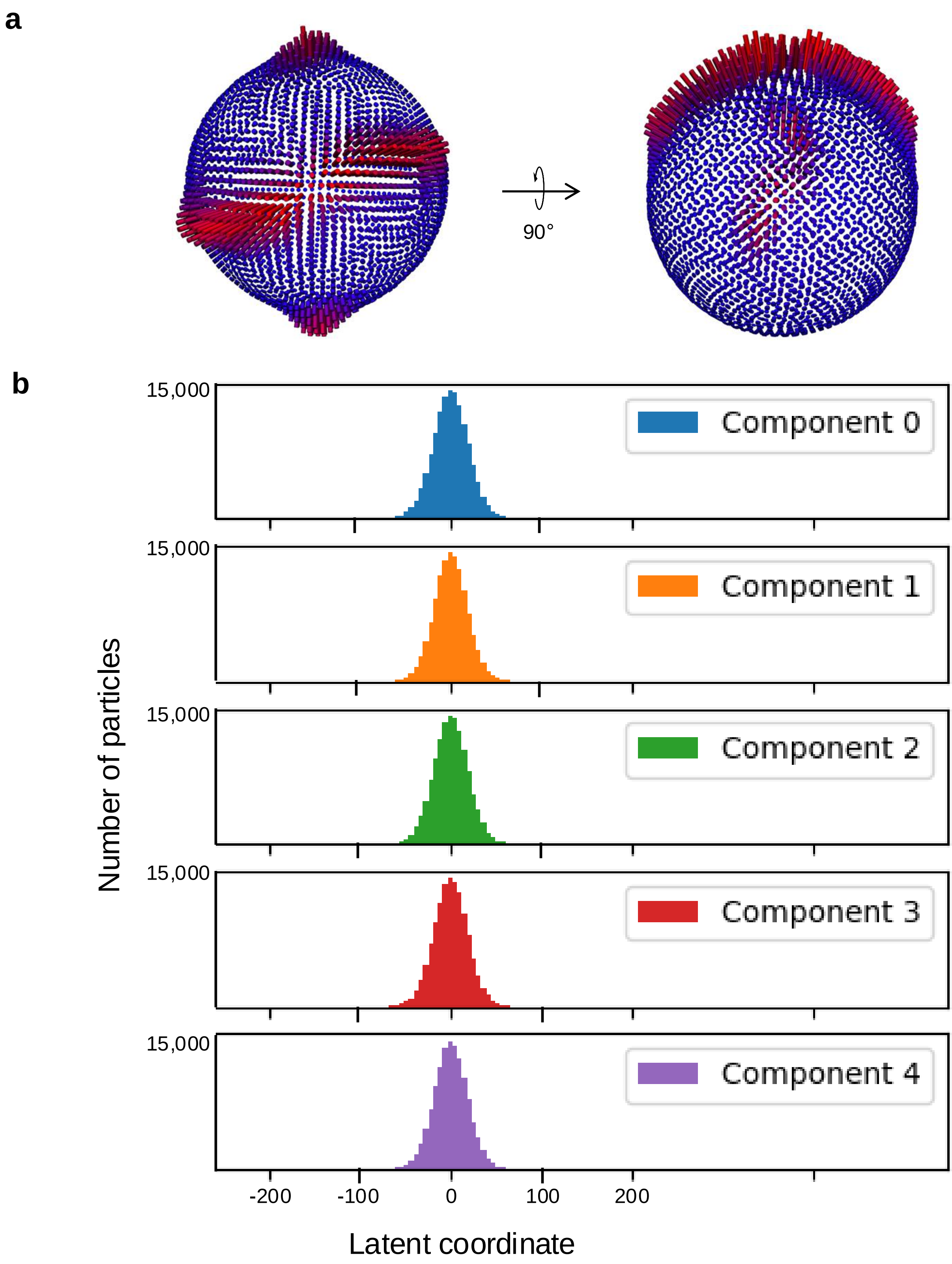

**Figure S31. PCA and particle distributions .**
**a** 3D representation of particle distribution for the reconstruction of ASNS R142I variant; 2 views at 90° are represented. Red cylinders indicate the amount of particle in each rotation angle for the reconstruction. **b** Particle distribution in latent space, represented per principal component as a function of latent coordinate.

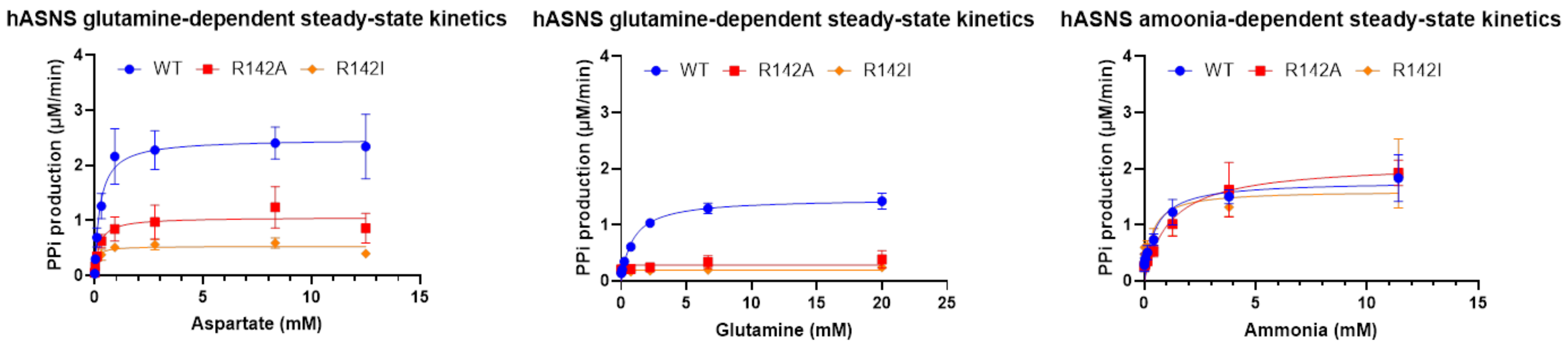

**Figure S32**. **Kinetic plots for the steady-state experiments.** (Left) Pyrophosphate production vs. L-**aspartate** concentration (0-12.5 mM), at fixed concentrations of L-glutamine (20 mM), 10 mM $MgCl_2$, 2 mM dithiothreitol and 5 mM ATP in 100 mM EPPS, pH 8 for WT ASNS and the R142I and R142A ASNS variants. (Middle) Pyrophosphate production vs. L-glutamine concentration (0-20 mM), at fixed concentrations of **aspartate** (10 mM), 10 mM $MgCl_2$, 2 mM dithiothreitol and 5 mM ATP in 100 mM EPPS, pH 8 for WT ASNS and the R142I and R142A ASNS variants. (Right) Pyrophosphate production vs. ammonia concentration (0-11.4 mM), at fixed concentrations of L-aspartate (10 mM), 10 mM $MgCl_2$, 2 mM dithiothreitol and 5 mM ATP in 100 mM EPPS, pH 8 for WT ASNS and the R142I and R142A ASNS variants. Data analysis was performed using GraphPad Prism.

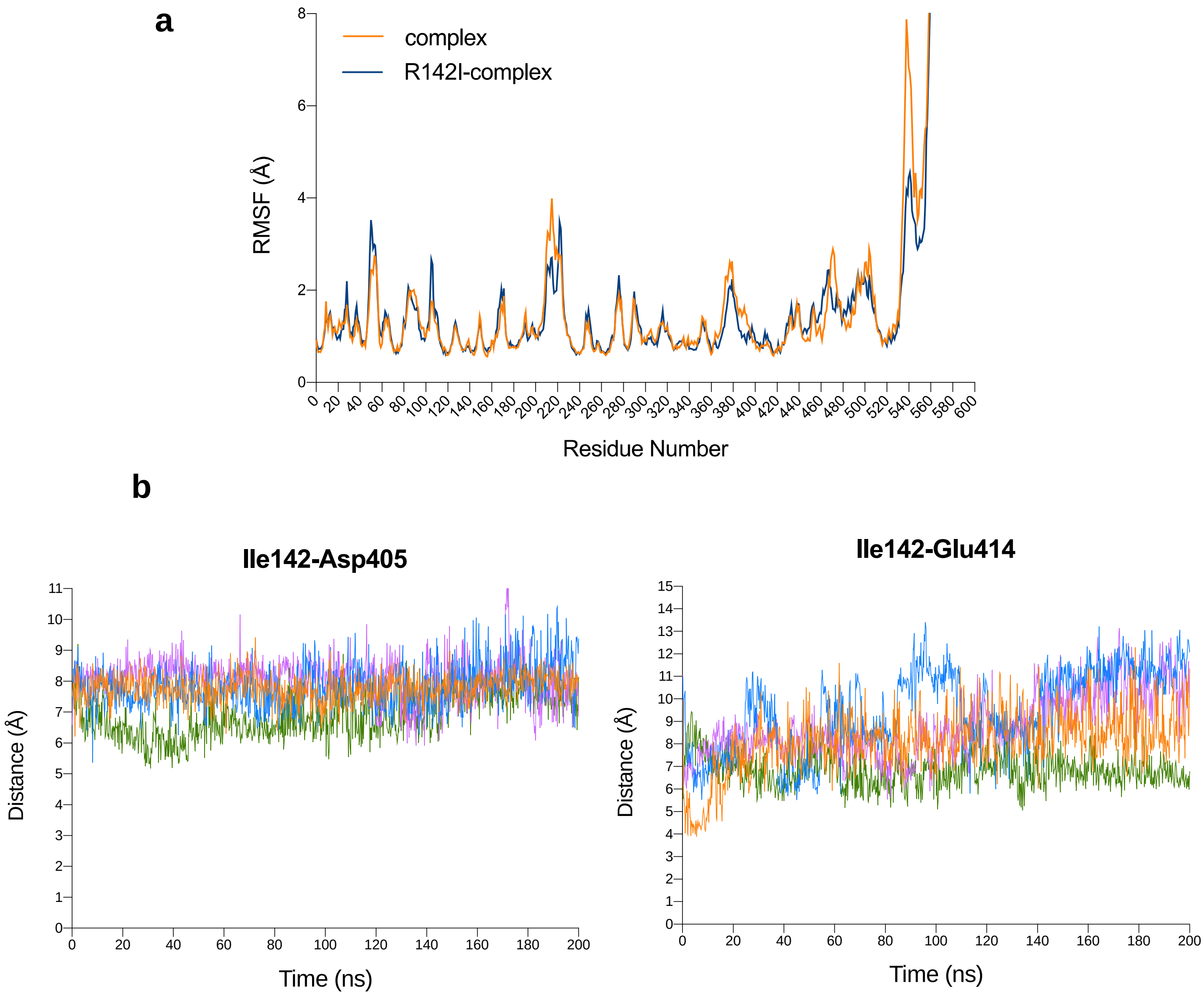

**Figure S33**. **200 ns MD simulations of the R142I/β-aspartyl-AMP/$PP_i$ ternary complex**. **a** Comparison of backbone RMSF values computed for the WT ASNS/β-aspartyl-AMP/$PP_i$ ternary complex (orange) and the R142I/β-aspartyl-AMP/$PP_i$ ternary complex (blue). **b** Replicate 200 ns MD simulations of the R142I/β-aspartyl-AMP/$PP_i$ ternary complex. (left) Plots of the distance between Ile-142 and Asp-405 and (right) between Ile-142 and Glu-414 in the four MD simulation replicates. Distances were calculated between the delta carbon atom of Ile-142 and and one of the carboxylate oxygen atoms of Asp-405 or of Glu-414, respectively. See Fig. S17 for the location of these oxygen atoms.

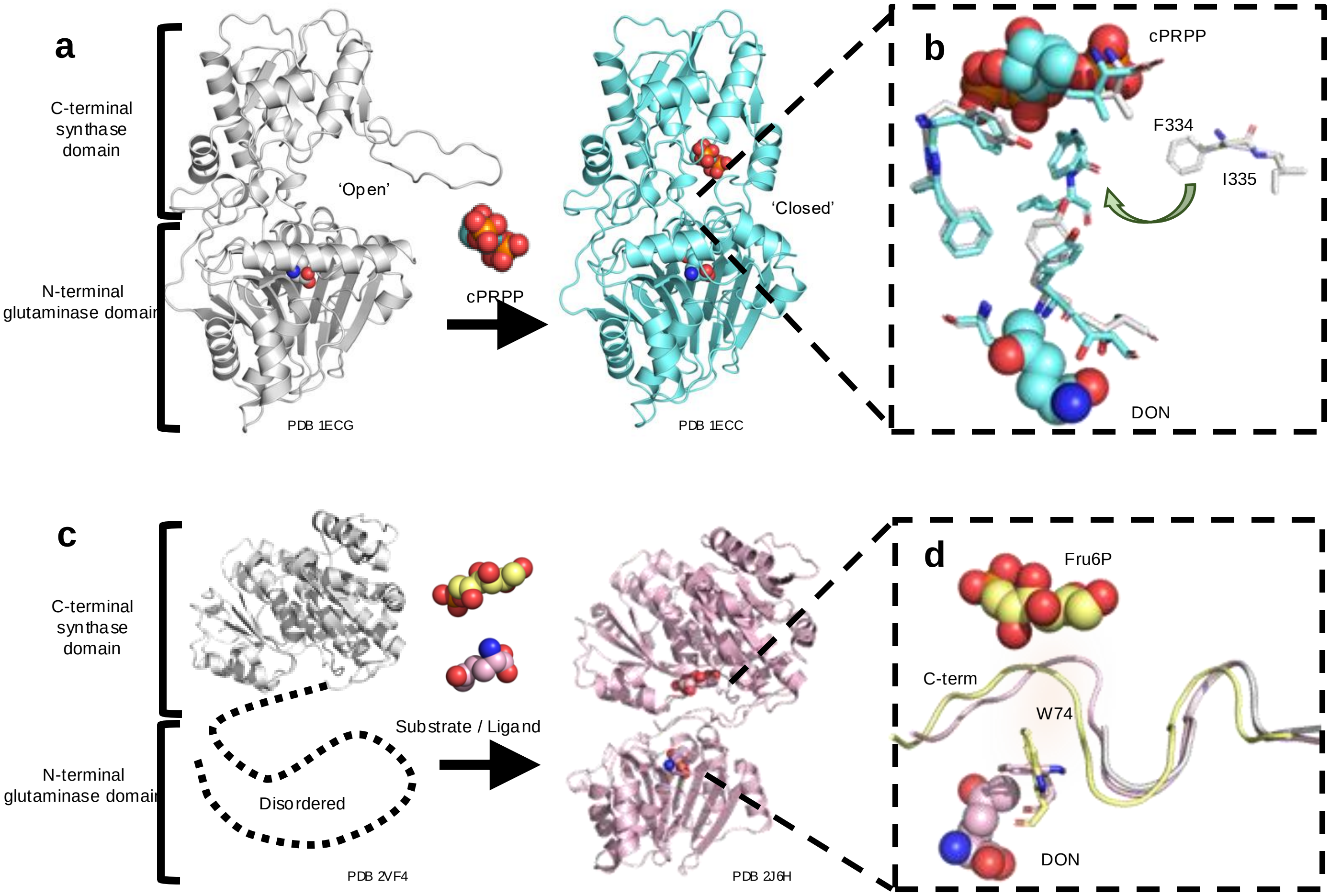

**Figure S34**. **a** cPRPP binding induces a conformational change in an active site loop of the synthase domain of glutamine PRPP amidotransferase (GPA). Cartoon representations of the apo-enzyme (PDB 1ECG) and the complex between cPRPP and DON-modified GPA (PDB 1ECC) are shown in gray and cyan, respectively. **b** When a stable PRPP analog (cPRPP) binds to the DON-modified GPA, the loop residues Phe-334 and Ile-335 reorient (indicated by the green arrow) to create a tunnel connecting the N- and C-terminal active sites (green shading). Residues in the apo-enzyme (PDB 1ECG) and the cPRPP/DON-modified GPA complex (PDB 1ECC) are shown as gray and cyan sticks, respectively. **c** Substrate/ligand binding is needed to stabilize the N-terminal domain of GFAT, which is not resolved in the X-ray crystal structure of the free enzyme (gray) (PDB 2VF4)[21]. When glucose-6-phosphate is present in DON-modified GFAT (light pink), however, both domains of the enzyme are observed (PDB 2J6H). Glucose-6-phosphate and DON are rendered as yellow and light pink spheres, respectively. **d** Close-up of the open intramolecular tunnel in the fructose-6-phosphate/DON-modified GFAT complex (yellow) (PDB 4AMV) formed by rotation of Trp-74. The C-terminal tail is disordered in the free enzyme (gray) but becomes ordered in the two complexes (pink, yellow). Fructose-6-phosphate and DON are rendered as yellow and light pink spheres, respectively.

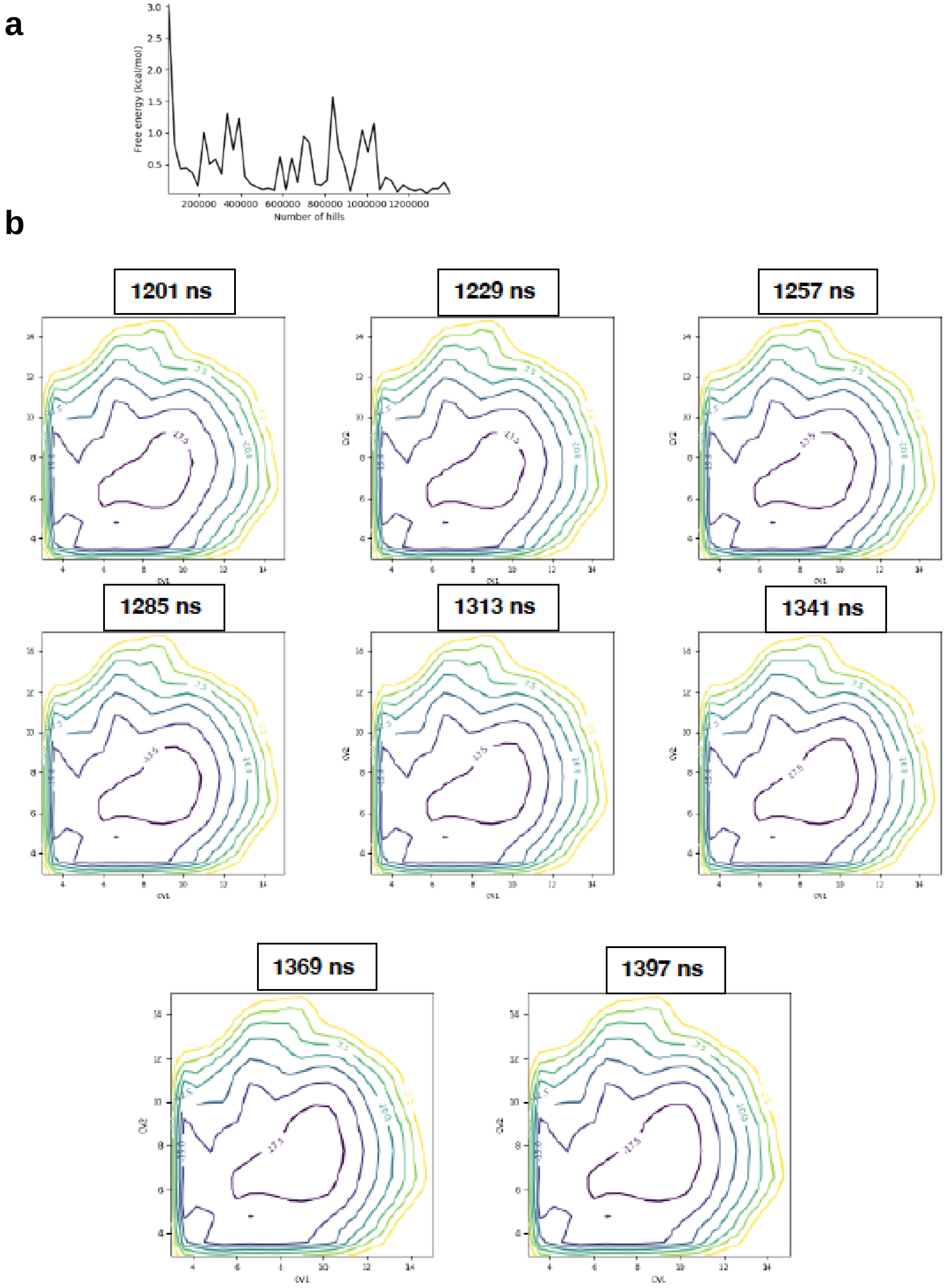

**Figure S35. Convergence of the metadynamics-based free energy surface for the Arg-142 side chain in the human apo-ASNS monomer.** Two distance-based collective variables were defined for these simulations (Supplementary Fig. S17), which correspond to the distances between the center-of-mass (COM) of the guanidinium group in Arg-142 and the COM of the carboxylate moieties in either Asp-405 (CV1) or Glu-414 (CV2). **a** Distance of free energies as a function of time, **b** Convergence of the free energy surface in the final stages of the MTD simulation.

**a**

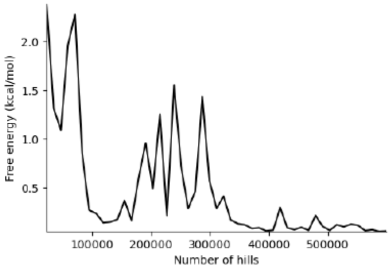

**b**

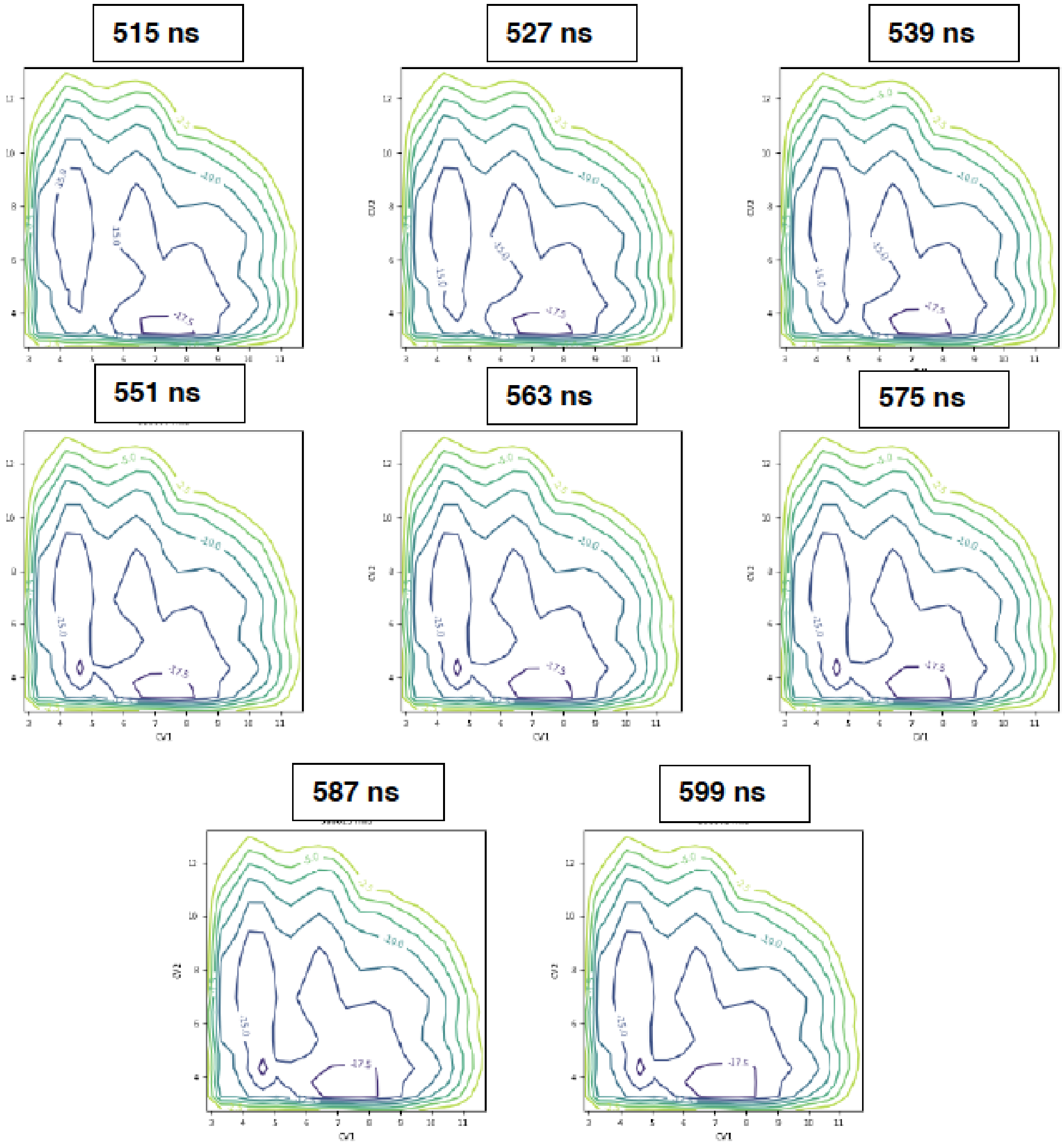

**Figure S36. Convergence of the metadynamics-based free energy surface for the Arg-142 side chain in the WT ASNS/β-aspartyl-AMP/$PP_i$ ternary complex.** Two distance-based collective variables were defined for these simulations (Supplementary Fig. S17), which correspond to the distances between the center-of-mass (COM) of the guanidinium group in Arg-142 and the COM of the carboxylate moieties in either Asp-405 (CV1) or Glu-414 (CV2). **a** Distance of free energies as a function of time, **b** Convergence of the free energy surface in the final stages of the MTD simulation.

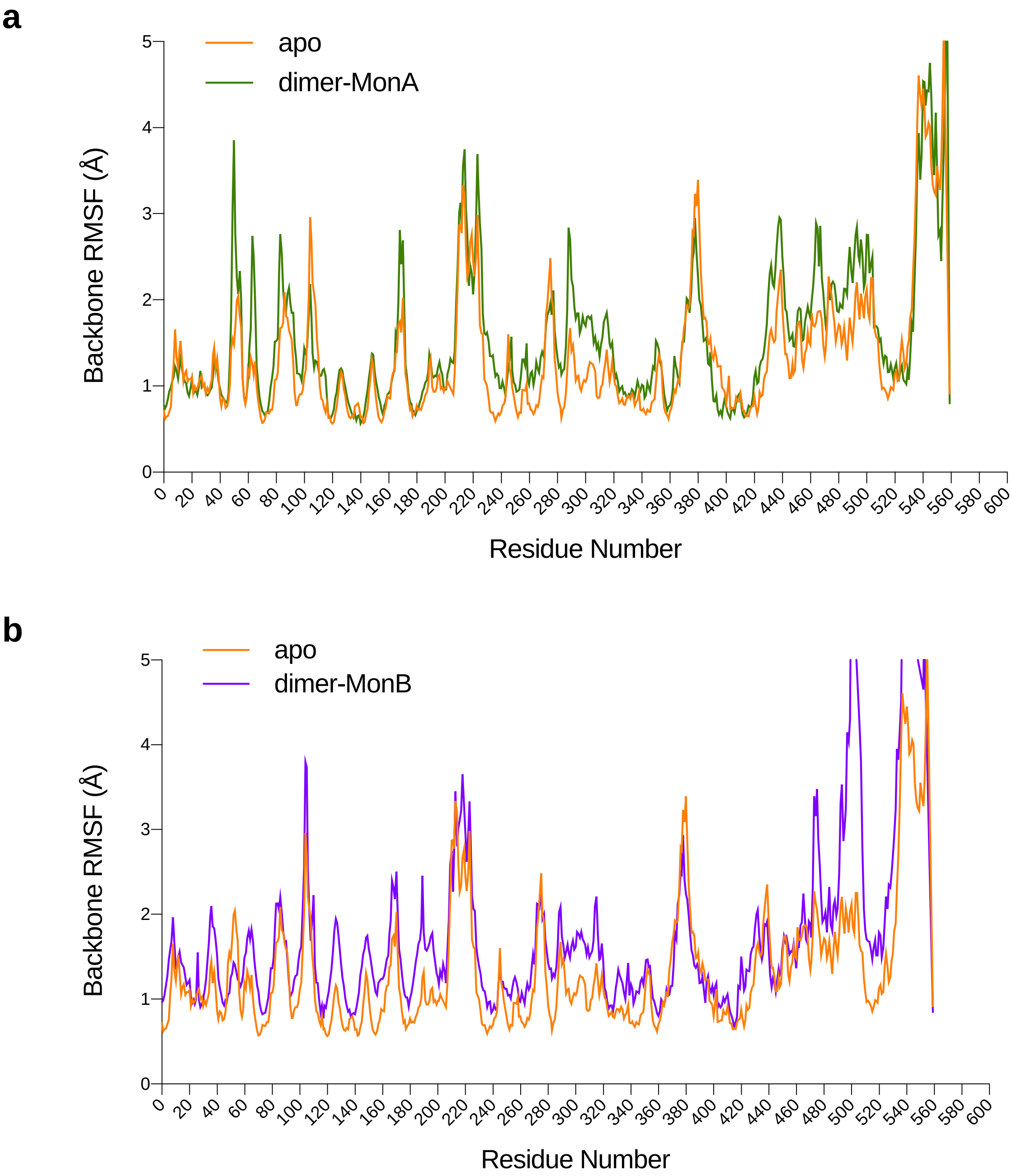

**Figure S37**. **Comparison of backbone RMSF values computed from 200 ns MD trajectories of the full-length apo-ASNS monomer and the apo-ASNS dimer. a** Values for the apo-ASNS monomer and monomer A in the dimer are shown in orange and green, respectively. **b** Values for the apo-ASNS monomer and monomer B in the dimer are shown in orange and purple, respectively. The deviations in the C-terminal region are not important for the tunnel structure and dynamics.

**Table S2:** Average values and standard deviation of nitrogen-oxygen distances (Å) between the Arg-142 side chain and selected residues, as calculated from four 200 ns replicate MD simulations of the human apo-ASNS monomer. Atom names correspond to the PDB naming convention for arginine, aspartate and glutamate residues. The molecular snapshots shown in Fig. 5a correspond to structures sampled from the trajectory of replica MD simulation 1.

| **WT ASNS** | Replica 1 | Replica 2 | Replica 3 | Replica 4 |
|---|---|---|---|---|
| **Asn-74** | | | | |
| OD1-NE | 4.5 ± 1.1 | 5.8 ± 1.1 | 6.1 ± 0.9 | 6.1 ± 2.3 |
| OD1-NH1 | 6.2 ± 1.0 | 6.1 ± 1.1 | 7.9 ± 0.9 | 8.0 ± 2.1 |
| OD1-NH2 | 6.2 ± 1.6 | 7.3 ± 1.1 | 8.0 ± 1.0 | 7.8 ± 2.3 |
| **Glu-76** | | | | |
| OE1-NE | 9.4 ± 1.0 | 7.7 ± 0.8 | 7.5 ± 1.6 | 6.5 ± 0.9 |
| OE2-NE | 9.5 ± 1.2 | 7.5 ± 0.8 | 8.1 ± 1.5 | 6.5 ± 1.0 |
| OE1-NH1 | 8.4 ± 0.9 | 7.7 ± 1.0 | 6.6 ± 1.8 | 7.4 ± 1.2 |
| OE2-NH1 | 8.5 ± 1.1 | 7.5 ± 1.0 | 7.6 ± 1.3 | 7.4 ± 1.4 |
| OE1-NH2 | 9.2 ± 1.3 | 6.2 ± 0.8 | 7.7 ± 1.5 | 5.9 ± 1.1 |
| OE2-NH2 | 9.3 ± 1.4 | 6.0 ± 0.8 | 8.6 ± 1.2 | 5.9 ± 1.1 |
| **Asp-405** | | | | |
| OD1-NE | 7.2 ± 0.9 | 6.5 ± 0.4 | 4.0 ± 0.3 | 7.2 ± 0.9 |
| OD2-NE | 6.5 ± 1.1 | 8.2 ± 0.4 | 5.1 ± 0.4 | 7.7 ± 0.9 |
| OD1-NH1 | 8.9 ± 1.3 | 5.4 ± 0.5 | 5.0 ± 0.3 | 7.6 ± 1.0 |
| OD2-NH1 | 8.2 ± 0.9 | 7.5 ± 0.5 | 6.6 ± 0.4 | 8.0 ± 0.9 |
| OD1-NH2 | 7.4 ± 1.1 | 7.4 ± 0.4 | 2.8 ± 0.4 | 8.5 ± 1.0 |
| OD2-NH2 | 6.6 ± 1.0 | 9.5 ± 0.4 | 4.5 ± 0.5 | 9.0 ± 1.0 |
| **Glu-414** | | | | |
| OE1-NE | 6.3 ± 1.0 | 4.8 ± 0.2 | 6.0 ± 1.0 | 4.6 ± 1.0 |
| OE2-NE | 6.7 ± 0.8 | 4.8 ± 0.2 | 6.7 ± 0.8 | 4.6 ± 1.0 |
| OE1-NH1 | 6.7 ± 1.4 | 2.8 ± 0.3 | 6.2 ± 1.1 | 5.3 ± 1.1 |
| OE2-NH1 | 7.1 ± 1.0 | 3.5 ± 0.4 | 7.3 ± 0.9 | 5.0 ± 1.1 |
| OE1-NH2 | 6.0 ± 1.6 | 3.5 ± 0.2 | 5.2 ± 1.1 | 3.9 ± 0.9 |
| OE2-NH2 | 6.4 ± 1.2 | 2.7 ± 0.2 | 6.3 ± 0.9 | 3.5 ± 0.1 |

**Table S3:** Shortest nitrogen-oxygen distances (Å) between the Arg-142 side chain and selected residues in the snapshots (Fig. 5a) obtained from the 200 ns MD simulation of the human apo-ASNS monomer. Atom names correspond to the PDB naming convention for arginine, aspartate and glutamate residues.

| **WT ASNS** | 20 ns | 80 ns | 140 ns | 200 ns |
|---|---|---|---|---|
| **Asn-74** | 2.7 (OD1-NH2) | 4.4 (OD1-NE) | 5.4 (OD1-NE) | 7.2 (OD1-NE) |
| **Glu-76** | 7.0 (OE1-NH2) | 7.1 (OE1-NH1) | 8.7 (OE1-NH1) | 7.1 (OE1-NH1) |
| **Asp-405** | 5.7 (OD1-NH2) | 6.9 (OD1-NH2) | 5.7 (OD2-NE) | 5.4 (OD2-NE) |
| **Glu-414** | 2.5 (OE1-NH2) | 7.2 (OE1-NH2) | 5.5 (OE1-NH2) | 5.0 (OE1-NE) |

**Table S4:** Average values and standard deviation of nitrogen-oxygen distances (Å) between the Arg-142 side chain and selected residues, as calculated from four 200 ns replicate MD simulations of the the human ASNS/β-aspartyl-AMP/MgPP$_i$ ternary complex. Atom names correspond to the PDB naming convention for arginine, aspartate and glutamate residues. The molecular snapshots shown in Fig. 5b correspond to structures sampled from the trajectory of replica MD simulation 1.

| Complex | Replica 1 | Replica 2 | Replica 3 | Replica 4 |
|---|---|---|---|---|
| **Asn-74** | | | | |
| OD1-NE | 4.8 ± 1.7 | 5.9 ± 1.8 | 5.2 ± 0.9 | 5.1 ± 0.9 |
| OD1-NH1 | 6.8 ± 1.5 | 8.0 ± 1.7 | 6.9 ± 1.1 | 6.8 ± 1.1 |
| OD1-NH2 | 6.5 ± 2.0 | 7.8 ± 2.0 | 6.5 ± 1.2 | 7.0 ± 0.9 |
| **Glu-76** | | | | |
| OE1-NE | 10.4 ± 1.4 | 8.2 ± 1.0 | 5.4 ± 1.5 | 6.9 ± 1.9 |
| OE2-NE | 10.4 ± 1.3 | 8.9 ± 1.3 | 5.5 ± 1.6 | 6.8 ± 1.9 |
| OE1-NH1 | 10.1 ± 1.4 | 10.0 ± 1.1 | 6.4 ± 1.5 | 8.0 ± 1.9 |
| OE2-NH1 | 10.1 ± 1.2 | 10.6 ± 1.5 | 6.6 ± 1.7 | 7.9 ± 2.0 |
| OE1-NH2 | 11.0 ± 2.0 | 8.3 ± 1.0 | 4.5 ± 1.8 | 6.1 ± 2.0 |
| OE2-NH2 | 11.0 ± 1.8 | 8.9 ± 1.4 | 4.7 ± 2.0 | 6.0 ± 2.0 |
| **Asp-405** | | | | |
| OD1-NE | 4.5 ± 0.7 | 4.7 ± 0.6 | 6.7 ± 1.1 | 4.1 ± 0.3 |
| OD2-NE | 5.3 ± 0.9 | 6.0 ± 0.4 | 8.2 ± 1.1 | 6.0 ± 0.3 |
| OD1-NH1 | 4.6 ± 1.4 | 4.9 ± 0.6 | 6.0 ± 1.3 | 3.0 ± 0.5 |
| OD2-NH1 | 5.6 ± 1.9 | 5.6 ± 0.3 | 7.4 ± 1.2 | 5.0 ± 0.4 |
| OD1-NH2 | 3.5 ± 1.3 | 5.9 ± 0.6 | 8.0 ± 1.3 | 4.9 ± 0.4 |
| OD2-NH2 | 4.3 ± 1.8 | 6.9 ± 0.5 | 9.3 ± 1.4 | 7.0 ± 0.3 |
| **Glu-414** | | | | |
| OE1-NE | 5.7 ± 1.1 | 3.7 ± 0.8 | 5.3 ± 0.8 | 6.3 ± 1.5 |
| OE2-NE | 5.9 ± 1.0 | 4.3 ± 0.6 | 5.8 ± 1.3 | 6.2 ± 1.5 |
| OE1-NH1 | 6.0 ± 1.1 | 5.0 ± 0.7 | 5.1 ± 1.1 | 6.6 ± 1.4 |
| OE2-NH1 | 6.3 ± 0.8 | 5.9 ± 0.7 | 5.7 ± 2.1 | 6.6 ± 1.3 |
| OE1-NH2 | 5.3 ± 1.4 | 3.1 ± 0.8 | 5.0 ± 1.4 | 5.3 ± 1.6 |
| OE2-NH2 | 5.7 ± 1.0 | 4.1 ± 0.9 | 5.4 ± 2.4 | 5.3 ± 1.6 |

**Table S5:** Shortest nitrogen-oxygen distances (Å) between the Arg-142 side chain and selected residues in the snapshots (Fig. 5b) obtained from the 200 ns MD simulation of the ASNS/β-aspartyl-AMP/$MgPP_i$ ternary complex. Atom names correspond to the PDB naming convention for arginine, aspartate and glutamate residues.

| **Complex** | 20 ns | 80 ns | 140 ns | 200 ns |
|---|---|---|---|---|
| **Asn-74** | 7.4 (OD1-NE) | 6.2 (OD1-NE) | 4.4 (OD1-NE) | 3.3 (OD1-NH2) |
| **Glu-76** | 9.1 (OE2-NH1) | 10.1 (OE2-NH1) | 9.0 (OE1-NH1) | 7.9 (OE2-NH2) |
| **Asp-405** | 5.0 (OD1-NE) | 2.7 (OD1-NH2) | 2.7 (OD2-NH2) | 2.6 (OD1-NH1) |
| **Glu-414** | 2.7 (OE1-NH2) | 4.6 (OE1-NE) | 5.2 (OE2-NH2) | 3.9 (OE2-NH2) |

**Table S6 List of Supplementary Movies.**

| Movie | | Content |
|---|---|---|
| M1 | | WT_3DVA_Component_0 |
| M2 | | WT_3DVA_Component_1 |
| M3 | | WT_3DVA_Component_2 |
| M4 | | WT_3DVA_Component_3 |
| M5 | | WT_3DVA_Component_4 |
| M6 | | R142I_3DVA_Component_0 |
| M7 | | R142I_3DVA_Component_1 |
| M8 | | R142I_3DVA_Component_2 |
| M9 | | R142I_3DVA_Component_3 |
| M10 | | R142I_3DVA_Component_4 |
| M11 | | MD_Trajectory_apo_WT_ASNS |
| M12 | | MD_Trajectory_ASNS_Ternary_Complex |
| M13 | | MD_Trajectory_apo_R142I_Variant |
| M14 | | MD_Trajectory_R142I_Ternary_Complex |

### Table S7. Ionization of residues using PROPKA

| | WT | R142A | R142I |
|---|---|---|---|
| CYS_1 | -1 | -1 | -1 |
| ASP_10 | -1 | -1 | -1 |
| ASP_11 | -1 | -1 | -1 |
| LYS_22 | 1 | 1 | 1 |
| HIS_25 | 0 | 0 | 0 |
| ARG_26 | 1 | 1 | 1 |
| ASP_29 | -1 | -1 | -1 |
| ARG_32 | 1 | 1 | 1 |
| GLU_34 | -1 | -1 | -1 |
| HIS_47 | 0 | 0 | 0 |
| ARG_48 | 1 | 1 | 1 |
| ASP_53 | -1 | -1 | -1 |
| ARG_62 | 1 | 1 | 1 |
| LYS_64 | 1 | 1 | 1 |
| LYS_65 | 1 | 1 | 1 |
| GLU_76 | -1 | -1 | -1 |
| HIS_80 | 0 | 0 | 0 |
| LYS_81 | 1 | 1 | 1 |
| LYS_82 | 1 | 1 | 1 |
| HIS_86 | 0 | 0 | 0 |
| GLU_88 | -1 | -1 | -1 |
| GLU_90 | -1 | -1 | -1 |
| LYS_94 | 1 | 1 | 1 |
| ASP_96 | -1 | -1 | -1 |
| GLU_98 | -1 | -1 | -1 |
| HIS_102 | 0 | 0 | 0 |
| ASP_105 | -1 | -1 | -1 |
| LYS_106 | 1 | 1 | 1 |
| GLU_110 | -1 | -1 | -1 |
| ASP_117 | -1 | -1 | -1 |
| ASP_126 | -1 | -1 | -1 |
| LYS_130 | 1 | 1 | 1 |
| LYS_131 | 1 | 1 | 1 |
| ARG_136 | 1 | 1 | 1 |
| ASP_137 | -1 | -1 | -1 |
| ARG_142 | 1 | - | - |
| LYS_146 | 1 | 1 | 1 |
| GLU_150 | -1 | -1 | -1 |
| ASP_151 | -1 | -1 | -1 |
| GLU_159 | -1 | -1 | -1 |
| LYS_161 | 1 | 1 | 1 |
| LYS_167 | 1 | 1 | 1 |
| HIS_168 | 0 | 0 | 0 |
| LYS_175 | 1 | 1 | 1 |
| GLU_177 | -1 | -1 | -1 |
| HIS_183 | 0 | 0 | 0 |
| GLU_185 | -1 | -1 | -1 |
| ASP_188 | -1 | -1 | -1 |
| LYS_190 | 1 | 1 | 1 |
| LYS_194 | 1 | 1 | 1 |
| GLU_199 | -1 | -1 | -1 |
| LYS_202 | 1 | 1 | 1 |
| HIS_204 | 0 | 0 | 0 |
| HIS_205 | 0 | 0 | 0 |
| ARG_207 | 1 | 1 | 1 |
| ASP_208 | -1 | -1 | -1 |
| HIS_212 | 0 | 0 | 0 |
| ASP_216 | -1 | -1 | -1 |
| GLU_219 | -1 | -1 | -1 |
| LYS_220 | 1 | 1 | 1 |
| GLU_226 | -1 | -1 | -1 |
| GLU_228 | -1 | -1 | -1 |
| LYS_231 | 1 | 1 | 1 |
| ARG_235 | 1 | 1 | 1 |
| LYS_243 | 1 | 1 | 1 |
| LYS_244 | 1 | 1 | 1 |
| ARG_245 | 1 | 1 | 1 |
| ASP_249 | -1 | -1 | -1 |
| ARG_250 | 1 | 1 | 1 |
| ARG_251 | 1 | 1 | 1 |
| ASP_261 | -1 | -1 | -1 |
| LYS_271 | 1 | 1 | 1 |
| LYS_274 | 1 | 1 | 1 |
| GLU_275 | -1 | -1 | -1 |

**Table S7. Ionization of residues using PROPKA**

| | WT | R142A | R142I |
|---|---|---|---|
| GLU_290 | -1 | -1 | -1 |
| ASP_291 | -1 | -1 | -1 |
| ASP_294 | -1 | -1 | -1 |
| ARG_299 | 1 | 1 | 1 |
| LYS_300 | 1 | 1 | 1 |
| ASP_303 | -1 | -1 | -1 |
| HIS_304 | 0 | 0 | 0 |
| GLU_308 | -1 | -1 | -1 |
| HIS_309 | 0 | 0 | 0 |
| GLU_311 | -1 | -1 | -1 |
| GLU_317 | -1 | -1 | -1 |
| GLU_318 | -1 | -1 | -1 |
| ASP_324 | -1 | -1 | -1 |
| GLU_325 | -1 | -1 | -1 |
| GLU_331 | -1 | -1 | -1 |
| ASP_334 | -1 | -1 | -1 |
| ARG_339 | 1 | 1 | 1 |
| LYS_349 | 1 | 1 | 1 |
| ARG_352 | 1 | 1 | 1 |
| LYS_353 | 1 | 1 | 1 |
| ASP_356 | -1 | -1 | -1 |
| GLU_364 | -1 | -1 | -1 |
| ASP_367 | -1 | -1 | -1 |
| GLU_368 | -1 | -1 | -1 |
| HIS_377 | 0 | 0 | 0 |
| LYS_378 | 1 | 1 | 1 |
| GLU_383 | -1 | -1 | -1 |
| LYS_384 | 1 | 1 | 1 |
| GLU_386 | -1 | -1 | -1 |
| GLU_387 | -1 | -1 | -1 |
| GLU_388 | -1 | -1 | -1 |
| GLU_390 | -1 | -1 | -1 |
| ARG_391 | 1 | 1 | 1 |
| ARG_394 | 1 | 1 | 1 |
| GLU_395 | -1 | -1 | -1 |
| ASP_400 | -1 | -1 | -1 |
| ARG_403 | 1 | 1 | 1 |

| | WT | R142A | R142I |
|---|---|---|---|
| ASP_405 | -1 | -1 | -1 |
| ARG_406 | 1 | 1 | 1 |
| HIS_411 | 0 | 0 | 0 |
| GLU_414 | -1 | -1 | -1 |
| ARG_416 | 1 | 1 | 1 |
| ASP_421 | -1 | -1 | -1 |
| HIS_422 | 0 | 0 | 0 |
| ARG_423 | 1 | 1 | 1 |
| GLU_434 | -1 | -1 | -1 |
| ARG_436 | 1 | 1 | 1 |
| LYS_439 | 1 | 1 | 1 |
| GLU_443 | -1 | -1 | -1 |
| LYS_444 | 1 | 1 | 1 |
| HIS_445 | 0 | 0 | 0 |
| ARG_448 | 1 | 1 | 1 |
| GLU_449 | -1 | -1 | -1 |
| GLU_452 | -1 | -1 | -1 |
| ASP_453 | -1 | -1 | -1 |
| LYS_459 | 1 | 1 | 1 |
| GLU_460 | -1 | -1 | -1 |
| ARG_464 | 1 | 1 | 1 |
| LYS_466 | 1 | 1 | 1 |
| GLU_467 | -1 | -1 | -1 |
| ASP_471 | -1 | -1 | -1 |
| LYS_477 | 1 | 1 | 1 |
| LYS_482 | 1 | 1 | 1 |
| GLU_486 | -1 | -1 | -1 |
| GLU_489 | -1 | -1 | -1 |
| HIS_490 | 0 | 0 | 0 |
| ASP_493 | -1 | -1 | -1 |
| ASP_494 | -1 | -1 | -1 |
| LYS_503 | 1 | 1 | 1 |
| LYS_510 | 1 | 1 | 1 |
| LYS_512 | 1 | 1 | 1 |
| GLU_513 | -1 | -1 | -1 |
| ARG_518 | 1 | 1 | 1 |
| GLU_522 | -1 | -1 | -1 |

**Table S7 (continued). Ionization of residues using PROPKA**

| | WT | R142A | R142I |
|---|---|---|---|
| ARG_523 | 1 | 1 | 1 |
| HIS_524 | 0 | 0 | 0 |
| ARG_528 | 1 | 1 | 1 |
| ASP_530 | -1 | -1 | -1 |
| HIS_534 | 1 | 1 | 1 |
| LYS_539 | 1 | 1 | 1 |
| ASP_545 | -1 | -1 | -1 |
| ARG_549 | 1 | 1 | 1 |
| HIS_553 | 0 | 0 | 0 |
| LYS_555 | 1 | 1 | 1 |
| LYS_559 | 1 | 1 | 1 |
